# Spontaneous cell morphogenesis via self-propelled actin filaments

**DOI:** 10.1101/2024.02.28.582450

**Authors:** Kio Yagami, Kentarou Baba, Takunori Minegishi, Shinji Misu, Hiroko Katsuno-Kambe, Kazunori Okano, Yuichi Sakumura, Yoichiroh Hosokawa, Naoyuki Inagaki

## Abstract

Cells frequently undergo spontaneous morphogenesis, yet the underlying mechanisms remain incompletely understood. While actin filaments are central to cell morphogenesis and are typically regulated by biochemical signaling, cells can form protrusions even without clear external cues, suggesting the existence of intrinsic physical mechanisms. Here, we report that actin filaments undergo directional movement driven by treadmilling, an ATP-fueled polymerization-disassembly cycle intrinsic to actin. These Self-propelled Treadmilling Actin filaments (SpTAs), exhibit stochastic yet directional motion, in a manner similar to self-propelled “particles” rather than the previously reported reaction-diffusion “waves”. SpTA arrival at the cell periphery drives membrane protrusion by orienting their polymerizing ends outwards. This SpTA accumulation, guided by nascent membrane curvature, further amplifies protrusion growth and expansion, driving cellular polarization for migration. Our findings establish actin filament as a novel class of active particle, providing a fundamental physical framework for understanding how molecular-scale motion leads to higher-order organization in living systems.

## Introduction

Biological systems exhibit complex shapes that undergo dynamic changes, crucial processes for their function. Among these, actin-based protrusions are essential for cellular morphogenesis and motility (1–3). These structures appear in various forms, such as lamellipodia and filopodia for cell migration (1–3), microvilli to enhance nutrient absorption (4, 5), and phagocytic cups to ingest microbes (3). Their formation has conventionally been explained by local polymerization and disassembly of actin filaments (F-actins), orchestrated by biochemical signaling that activates key regulator proteins, such as the Arp2/3 complex, formins and ADF/cofilin (1–3, 6–9). Notably, cells exhibit the remarkable ability to spontaneously extend protrusions, even in the absence of explicit signaling cues. This suggests the existence of an intrinsic non-equilibrium mechanism that drives these structural transformations beyond localized signaling.

Actin filaments undergo wave-like propagation, termed actin “waves”, in various cell types (10–13). These phenomena have primarily been modeled by the reaction-diffusion or Turing-type frameworks, involving positive and negative feedback interactions (10–13). However, the underlying molecular basis and physical principles that validate these propagation models remain unclear (13). In parallel, the field of active matter physics has seen a surge of interest in self-propelled particles (also known as self-propelled Brownian particles, active particles or microswimmers) (14–17). They can take energy from their environment and convert it into directed motion to propel themselves, and their trajectories involve random fluctuations. Recent computational simulations and microswimmer studies have shown that these particles, when confined within cell-like vesicles, can spontaneously induce diverse membrane deformations remarkably similar to those observed during cellular morphogenesis (18–21).

F-actins have the inherent property to undergo energy-driven directional polymerization-disassembly cycle, known as treadmilling (1, 2, 9, 22). In this process, ATP-bound actin monomers polymerize at the plus ends of F-actins. This is followed by ATP hydrolysis within the filaments, which in turn leads to F-actin disassembly from the minus ends as ADP-bound actin molecules destabilize F-actins. Thus, F-actins undergo treadmilling by consuming ATP. Here we show that F-actin assemblies, which we term

Self-propelled Treadmilling Actin filaments (SpTAs), emerge widely in cells and translocate by treadmilling. Crucially, unlike to the continuous propagation of diffusion-reaction based actin wave, SpTAs exhibit random change in their direction of movement and accumulate at protrusive plasma membrane regions in a manner similar to the self-propelled particles (14). We propose that SpTAs broaden our understanding of intracellular actin dynamics and illuminate previously unresolved aspects of cellular behavior, including spontaneous cell morphogenesis. Indeed, perturbing SpTA movement impairs the spontaneous cell polarization essential for migration. Therefore, F-actins as “particles” rather than “waves” bridges the conceptual gap between active matter physics and cytoskeletal dynamics, providing a new framework for illustrating how molecular-scale motion leads to higher-order organization in living systems.

## RESULTS

### F-actin assemblies emerge widely in cells and translocate randomly

To investigate the intracellular actin dynamics in detail, we analyzed a subline of U251 glioma cells that spontaneously polarize and migrate in the absence of local signaling cues (Fig. S1*A*) (23). These cells actively form two types of protrusions, flattened lamellipodia and thin filopodia, at the front (Fig. S1*B*). Actin dynamics near the ventral plasma membrane were visualized with the F-actin marker LifeAct using total internal reflection fluorescence (TIRF) microscopy (Fig. 1*A*). Notably, assemblies of F-actins that form filopodium-like and lamellipodium-like structures emerged widely near the ventral plasma membrane, moved in random directions, and disappeared (Movie S1). The majority of them were filopodium-like (arrowheads, Fig. 1*B* and Movie S1 right) and randomly changed their direction of translocation (yellow arrowheads, Fig. 1*B* and Movie S1 right). The filopodium-like and lamellipodium-like F-actin assemblies were interchangeable: some filopodium-like F-actin assemblies emerged (yellow and cyan arrowheads) from lamellipodium-like assemblies (arrows) or branched off (green arrowheads) from filopodium-like assemblies (Fig. 1*B* and Movie S1 right). We also observed lamellipodium-like assemblies (arrows, Fig. S1*C*, Movie S1 right) expanded from filopodium-like assemblies (arrowheads). Similar movement of F-actin assemblies were observed in COS7 cells (Fig. S1*D*, Movie S1), suggesting that they are not cell type specific.

**Fig. 1.**
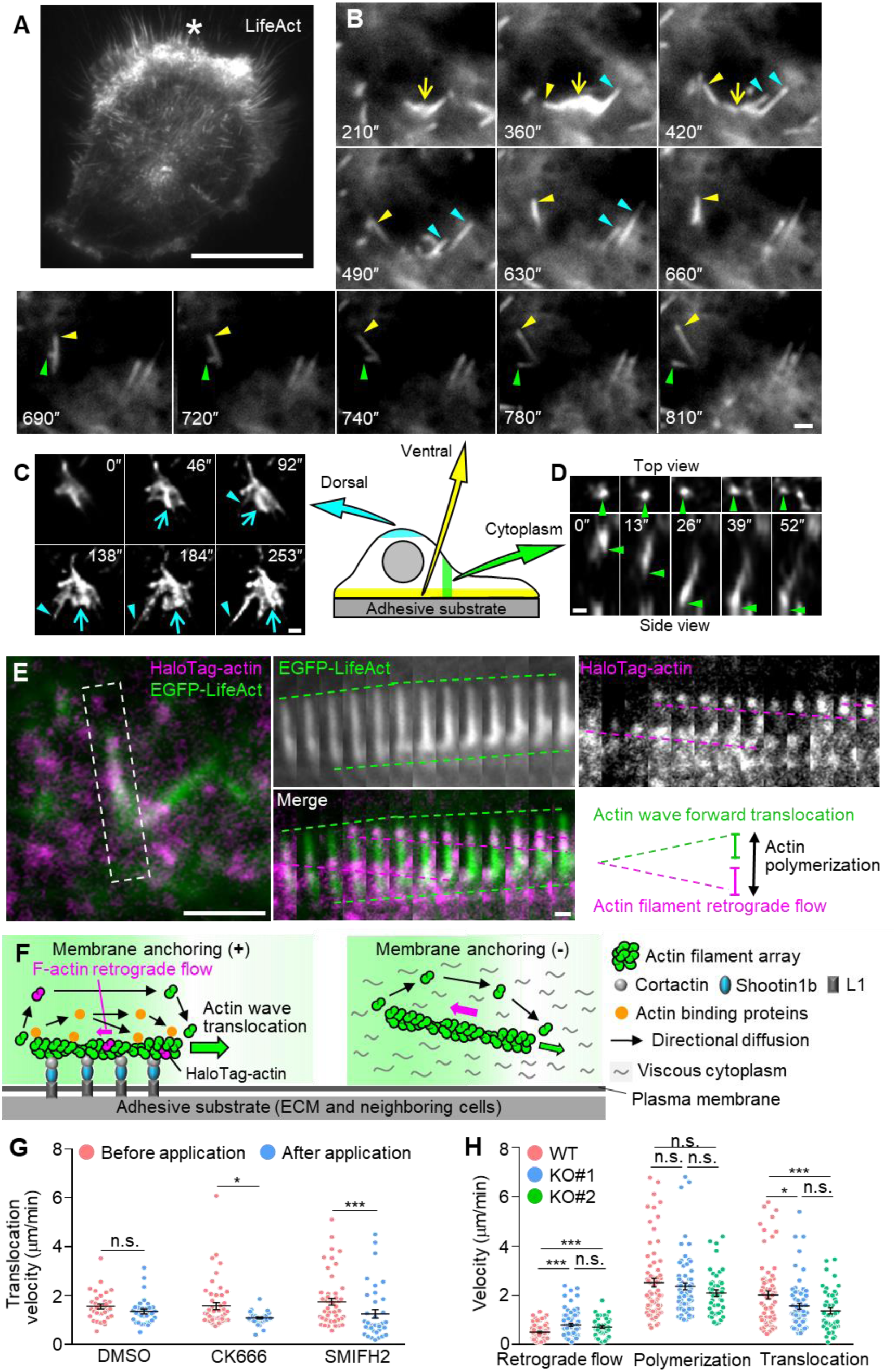
F-actin assemblies emerge widely in cells and translocate by treadmilling. (*A*) A fluorescence image of a U251 cell expressing EGFP-LifeAct obtained by TIRF microscopy. Asterisk indicates cellular leading edge. See Movie S1. Scale bar, 20 µm. (*B*) Fluorescence time-lapse images of filopodium-type (arrowheads) and lamellipodium-type (arrows) F-actin assemblies in U251 cells expressing LifeAct-mCherry obtained by TIRF microscopy. Filopodium-type F-actin assemblies frequently changed their direction of translocation (yellow arrowheads): some emerged and separated (yellow and cyan arrowheads) from lamellipodium-type assemblies (arrows) or branched off from filopodium-type assemblies (green arrowheads See Movie S1. Scale bar, 2 µm. (*C* and *D*) Fluorescence time-lapse images of F-actin assemblies on the dorsal membrane (blue arrowheads and arrows, C) and filopodium-type assemblies in cytoplasm (green arrowheads, D) in U251 cells expressing LifeAct-mCherry obtained by 3D imaging with confocal deconvolution microscopy. See Movie S2. Scale bars, 1 µm. (*E*) A fluorescent speckle image of HaloTag-actin in a U251 cell obtained by TIRF microscopy; F-actins were also monitored by EGFP-LifeAct. Time-lapse montages of a filopodium-type F-actin assembly in the rectangular region at 10-sec intervals are shown to the right. Green and magenta lines indicate forward translocation and actin filament retrograde flow, respectively. Actin polymerization rate can be calculated as the sum of the forward translocation rate and actin retrograde flow rate (double-headed arrow). See Movie S3. Scale bars, 5 µm (left); 1 µm (right). (*F*) A diagram showing the translocation mechanisms of the F-actin assembly travelling on the ventral plasma membrane (left). Right panel shows a possible mode of F-actin assembly translocation observed in Fig. 1C and *D*. For explanations, see text. (*G*) Translocation velocities of filopodium-type F-actin assemblies (green line, A) before and after the application of DMSO (control, N = 4 cells: before application, n = 34; after application, n = 32), CK666 (N = 3 cells: before application, n = 49; after application, n = 33) or SMIFH2 (N = 5 cells: before application, n = 52; after application, n = 37). Data represent means ± SEM; ***p < 0.01; *p < 0.05; ns, not significant. Statistical analyses were performed using the two-tailed Mann‒Whitney U-test. (*H*) Velocities of actin filament retrograde flow (magenta line, *E*), actin filament polymerization (double-headed arrow, *E*), and translocation (green line, *E*) of filopodium-type F-actin assemblies in WT and shootin1b KO U251 cells (WT, N = 11 cells, n = 71; KO#1, N = 9 cells, n = 69; KO#2, N = 10 cells, n = 45). Data represent means ± SEM; ***p < 0.01; *p < 0.05; ns, not significant. Statistical analyses were performed using the two-tailed Mann‒Whitney *U*-test. See also Figs. S1 and S2.

3D imaging with confocal deconvolution microscopy revealed F-actin assemblies moving along the dorsal plasma membrane (Fig. 1*C*) as well as those translocating through the cytoplasm without interacting with the plasma membrane (Fig. 1*D*, Movie S2). In addition, a combination of TIRF microscopy, which visualizes the region near the ventral plasma membrane, and epifluorescence microscopy, which visualizes the entire cell thickness, revealed a substantial fraction of F-actin assemblies translocating along the ventral plasma membrane (arrowheads, Fig. S1*E*, Movie S2), as well as those moving apart from the adhesive substrate at the leading edge (arrowheads, Fig. S1*F*, Movie S2). Thus, these F-actin assemblies emerge and translocate widely within cells, dynamically changing their shape and direction.

### Actin treadmilling drives translocation of F-actin assemblies

To analyze the mechanism that translocates F-actin assemblies, we monitored their subunit, actin molecules, by speckle imaging of HaloTag-actin (24). Actin molecules within the filopodium- and lamellipodium-type assemblies, advancing along the ventral membrane (Figs. 1*E* and S2*A*), moved retrogradely (magenta lines, Movie S3). Similar retrograde movement of actin molecules was observed in F-actin assemblies advancing without interaction with the plasma membrane (Fig. S2*B*), indicating that the F-actins in the forward-advancing F-actin assemblies undergo retrograde flow, accompanied by the polymerization at the front and disassembly at the rear: treadmilling (Fig. S2*C*).

We previously reported F-actin assemblies that translocate one-dimensionally along neuronal axons through treadmilling (25). Their front and rear ends polymerize and disassemble, respectively (black arrows, Fig. 1*F* left) (12, 25). In addition, they are weakly anchored to the plasma membrane and adhesive substrate, by the linker molecules shootin1a and cortactin and the cell adhesion molecule L1 (25, 26). As this anchoring impedes the retrograde flow (slippage) of treadmilling F-actins (magenta arrow, Fig. 1*F* left), the actin polymerization/disassembly rate exceeds the retrograde flow rate, resulting in the forward translocation of the front and rear ends of F-actins (green arrow). During the translocation, local concentrations of actin subunits and actin-binding proteins decrease at the front by polymerization and increase at the rear by disassembly, resulting in their diffusion along the local concentration gradients (black arrows, Fig. 1*F* left). Accompanied by this directional diffusion, treadmilling F-actin assemblies translocate actin and actin-binding proteins along axons (25, 27).

Immunoblot analyses showed that U251 cells express, shootin1b, cortactin and L1 (Fig. S2*D*); shootin1b is a splicing variant of shootin1a that also mediates the actin-L1 linkage (28). Shootin1b co-localized with the F-actin assemblies (arrowheads, Fig. S2*E*). Partial inhibition of actin polymerization by cytochalasin B at a low concentration (0.05 µM) (25) decreased the velocity of F-actin assembly translocation (Fig. S2*F*). The Arp2/3 complex inhibitor CK666 (100 µM) (29) or a relatively non-specific formin inhibitor SMIFH2 (5 µM) (30) also reduced the translocation velocity (Fig. 1*G*), indicating that their translocation is driven by actin polymerization. Furthermore, shootin1b knockout (KO) (Fig. S2*G*) increased the retrograde flow velocity of F-actins in the F-actin assemblies, without affecting the polymerization rate (Fig. 1*H*), indicating that shootin1b impedes the retrograde flow of treadmilling F-actins by anchoring them to the plasma membrane (for explanation see also Fig. S2*H*). Consistently, shootin1b KO also reduced the velocity of F-actin assembly translocation (Fig. 1*H*). Together, these results indicate that actin treadmilling drives translocation of F-actin assemblies and that shootin1b promotes their translocation by impeding their retrograde flow (Fig. 1*F* left).

We have also observed F-actin assemblies translocating apart from the adhesive substrate (Fig. 1*C* and Fig. S1*F*), as well as in the cytoplasm without interacting with the plasma membrane (Fig. 1*D*). Moreover, the inhibition of F-actin anchorage to the plasma membrane by shootin1b KO did not completely prevent their translocation (Fig. 1*H*). The cytoplasm represents a viscous and crowded intracellular environment containing proteins, macromolecules, and cytoskeletal components (31, 32). We consider that treadmilling F-actin assemblies can move forward without anchoring to the membrane when retrograde F-actin flow is mechanically obstructed in the intracellular environment (Fig. 1*F* right).

### Treadmilling F-actin assemblies move as self-propelled particles and form cell protrusions

When filopodium- and lamellipodium-type F-actin assemblies arrived at the cell periphery, they pushed the plasma membrane to form filopodia (arrowheads, Fig. 2*A*, Movie S4) and lamellipodia (yellow arrows, Fig. 2*B*, Movie S5). We also observed filopodium-type assemblies entering pre-existing lamellipodia (arrowheads, Fig. 2*C*, Movie S6 left) and filopodia (Movie S6 right). Their entrance in the lamellipodia resulted in local F-actin accumulation and lamellipodial expansion (arrows, Fig. 2*C*, Movie S6 left). Some of them continued to move laterally along the membrane (Fig. 2*A* and *B*). We also noted lamellipodium-type F-actin assemblies (yellow arrows, Fig. 2*B*, Movie S5) that merged with the pre-existing lamellipodia (cyan arrows) through their lateral movement.

**Fig. 2.**
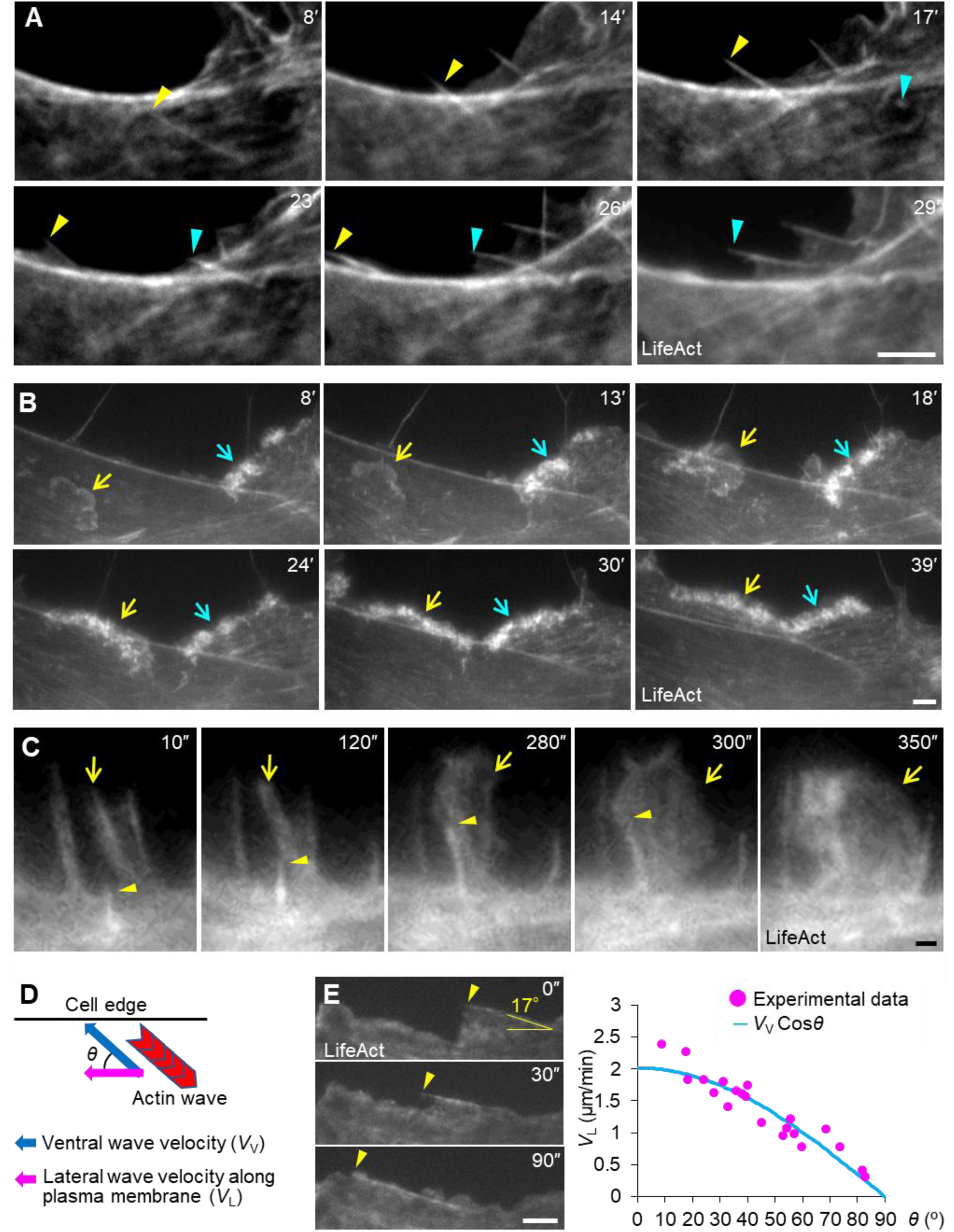
Treadmilling F-actin assemblies as precursors of protrusions. (*A* and *B*) Fluorescence time-lapse images of filopodium-type (*A*) and lamellipodium-type (*B*) F-actin assemblies arrived at the cell periphery in U251 cells expressing EGFP-LifeAct obtained by epifluorescence microscopy (*A*) and expressing LifeAct-mCherry obtained by TIRF microscopy (*B*). They pushed the plasma membrane, resulting in the formation of filopodia (arrowheads) and lamellipodium (yellow arrows), and continued to move laterally along the membrane. The latter merged with the pre-existing lamellipodium (blue arrows). See Movies S4 and S5. Scale bars, 5 µm. (*C*) Fluorescence time-lapse images of a filopodium-type F-actin assembly (arrowheads) which entered into a pre-existing lamellipodium (arrows) in U251 cells expressing EGFP-LifeAct obtained by TIRF microscopy. Its entry into lamellipodia resulted in local actin filament accumulation and lamellipodial extension (arrows). See Movie S6 (left). Scale bar, 1 µm. (*D*) A diagram describing the relationship between the velocities of lateral movement (VL) and ventral movement (VV) of filopodium-type F-actin assemblies. If F-actin assemblies move laterally along the membrane driven by directional actin treadmilling, VL is equal to VV cos*θ*, where *θ* is the angle of the polymerizing F-actins with respect to the membrane. (*E*) Fluorescence time-lapse images of a filopodium-type F-actin assembly travelling along the lateral membrane (arrowheads) of a U251 cell expressing LifeAct-mCherry obtained by epifluorescence microscopy. See Movie S7. The right graph shows VL and *θ* obtained by the time-lapse imaging; VL closely matched VVcos*θ* (N = 10 cells, n = 21). Scale bar, 5 µm.

If the lateral movements of filopodia and lamellipodia are propelled by treadmilling, their velocity (magenta arrow, Fig. 2*D*) would be related to the angle *θ* of their polymerizing F-actins with respect to the membrane and the average velocity of the treadmilling F-actin assemblies before arriving at the membrane (blue arrow). To examine this possibility, we focused on the motion of filopodium-type assemblies because the majority of the F-actin assemblies are the filopodium-type and filopodium- and lamellipodium-types are interchangeable (Movie S1). Time-lapse measurements of both the angle *θ* and the velocity of the filopodial lateral movement (*V*_L_) (Fig. 2*E*, Movie S7) demonstrated that the velocity closely matched the cosine *θ* for the average velocity of the ventrally translocating F-actin assemblies (*V*_V_) (2.0 ± 0.2 µm/min, Figs. 1*H* and 2*E*). Taken together, these data indicate that the treadmilling F-actin assemblies and filopodia/lamellipodia are equivalent structures and that they are propelled by actin treadmilling. Importantly, the actin filament assemblies move as discrete “particles” and do not propagate uniformly as “waves”. In addition, they randomly change the direction of movement and their arrival at the cell periphery results in local membrane protrusion, similar to the self-propelled particles (14). Thus, we refer to these actin assemblies as Self-propelled Treadmilling Actin filaments (SpTAs).

### SpTAs spontaneously accumulate at cell periphery and protrusions

We observed translocation of filopodium- and lamellipodium-type SpTAs in protrusive regions through their lateral movement (Fig. 3*A* and *B*, Movie S8), suggesting that their motion is affected by cell shape. To analyze a possible influence of cell shape on the SpTAs, we prepared a triangular pattern of adhesive laminin on the substrate (Fig. S3). U251 cells showed the triangular shape when cultured on the laminin-coated adhesive island, and F-actins accumulated at the corners of the triangular cells (Fig. 3*C*). Live imaging of U251 cells labelled with LifeAct-mCherry revealed the dynamic accumulation of F-actins at the corners (Fig. 3*D* and Movie S9). SpTAs travelled along the lateral edge, resulting in local accumulation of F-actins at the corners upon arrival (arrowheads). An Arp2/3 complex component, ARPC2, also accumulated at the protruding corners in the absence of local growth factor stimulation (Fig. 3*C*).

**Fig. 3.**
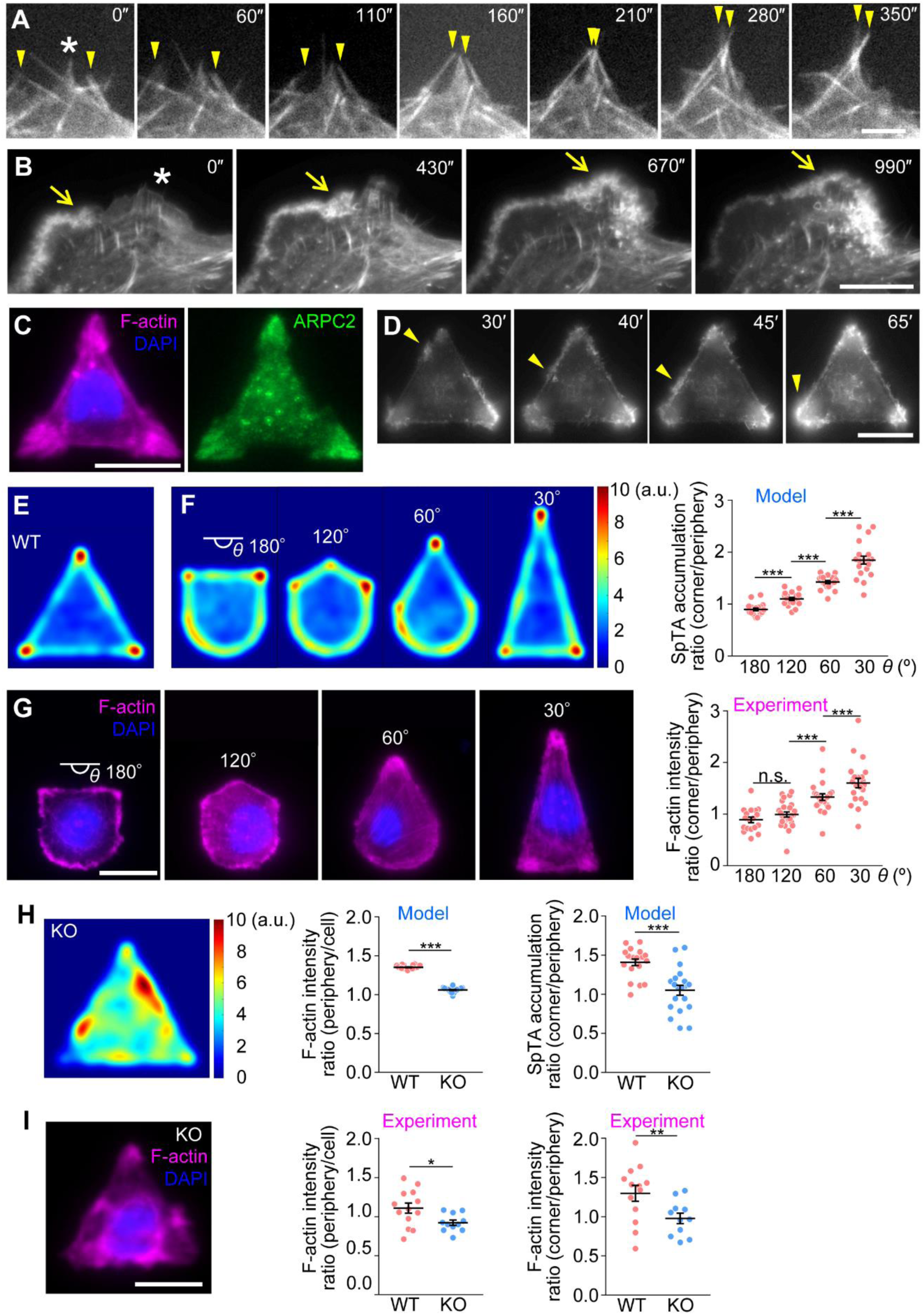
SpTAs mediate actin filament accumulation at the cell periphery and protrusions. (*A* and *B*) Fluorescence time-lapse images of filopodium-type (arrowheads, *A*) and lamellipodium-type (arrows, *B*) SpTAs which accumulated in protrusive regions (asterisks) in U251 cells expressing EGFP-LifeAct obtained by epifluorescence microscopy. See Movie S8. Scale bars, 20 µm (*B*); 5 µm (*A*). (*C*) A U251 cell cultured on a laminin-coated triangular adhesive island and stained with Alexa Fluor 594 conjugated phalloidin (for actin filament), anti-ARPC2 antibody, and DAPI (for nucleus). Scale bar, 20 µm. (*D*) Fluorescence time-lapse images of a U251 cell expressing LifeAct-mCherry cultured on a triangular adhesive island. The images were obtained by epifluorescence microscopy. Arrowheads indicate an SpTA travelled along the lateral membrane, resulting in a local accumulation of F-actins at the corner upon arrival. See Movie S9. Scale bar, 20 µm. (*E* and *F*) Mathematical model data showing SpTA localization in a triangular cell (*E*) and cells with different corner angles, 30°, 60°, 120° or 180° (*F*), presented in arbitrary unit (a.u.). See Movie S10. The right graph in (*F*) shows quantitative data for the SpTA accumulation at the corners with different corner angles (30-180°, n = 20 model data). Data represent means ± SEM; ***p < 0.01. Statistical analyses were performed using the two-tailed unpaired Student′s *t*-test (180° vs 120°, 120° vs 60°) and two-tailed unpaired Welch′s *t*-test (60° vs 30°). (*G*) U251 cells cultured on laminin-coated adhesive islands with the different corner angles and stained with Alexa Fluor 594 conjugated phalloidin and DAPI. The right graph shows quantitative data for the actin filament accumulation at the corners (30°, N = 6 experiments, n = 22 cells; 60°, N = 8 experiments, n = 23 cells; 120°, N = 7 experiments, n = 26 cells; 180°, N = 8 experiments, n = 18 cells). Scale bar, 20 µm. Data represent means ± SEM; ***p < 0.01; ns, not significant. Statistical analyses were performed using the two-tailed unpaired Student′s *t*-test (180° vs 120°) and two-tailed Mann‒Whitney *U*-test (120° vs 60°, 60° vs 30°). (*H*) Mathematical model data showing SpTA localization in a triangular shootin1b KO cell. See Movie S10. The right graphs show quantitative data for SpTA accumulation at the edge and corners of WT and shoootin1b KO cells (WT, KO, n = 20 model data). Data represent means ± SEM; ***p < 0.01. Statistical analyses were performed using the two-tailed unpaired Student′s *t*-test. (*I*) A shootin1b KO#1 U251 cell cultured on a laminin-coated triangular adhesive island and stained with Alexa Fluor 594 conjugated phalloidin and DAPI. The right graphs show quantitative data for actin filament accumulation at the edge and corners of WT and shoootin1b KO cells (WT, N = 3 experiments, n = 13 cells; KO#1, N = 6 experiments, n = 11 cells). Scale bar, 20 µm. Data represent means ± SEM; **p < 0.02; *p < 0.05.

Since the arrival of SpTAs increased F-actins at the protrusive areas (arrowheads, Fig. 3*D*), we hypothesized that they spontaneously accumulate at cell protrusions. To examine this possibility, we constructed a mathematical model to describe filopodium-type SpTAs, as we could observe F-actins clearly in these structures (Fig. 1*B*). Our model consists of the following properties of SpTA movement (see Materials and Methods for details).

1. SpTAs emerge at random locations within the cell and move in random directions (Movie S1).
2. SpTAs frequently change their direction of translocation and disappear at the end of their lifetime (Movie S1).
3. SpTAs re-emerge at the location where the preceding SpTA disappeared (see Materials and Methods).
4. SpTAs located at the cell periphery move laterally along the cell membrane. The velocity of the lateral movement conforms to the cos*θ* of the mean velocity of ventrally translocating SpTAs (*V*_V_), where *θ* is the angle of the direction of F-actins relative to the cell membrane (Fig. 2*E*).

All the model parameters, i.e., lifetime, length, moving velocity and degree of the change in moving direction, were obtained from our quantitative experimental data (Fig. S4*A*-*E*). The SpTAs described by the model accumulated dynamically at the corners of the triangular cells (Fig. 3*E* and Movie S10), as observed in U251 cells. Using this mathematical model, we further analyzed the movement of SpTAs in cells with different corner angles (i.e., 30°, 60°, 120° or 180°; Fig. S4*F*) and quantified the relative accumulation of F-actins at the cell periphery and corners (Fig. S4*G*). The SpTAs accumulated at the cell periphery (Fig. S4*H*). Furthermore, the accumulation of SpTAs at the corner increased as the corner angle decreased from 180° to 30° (Fig. 3*F*). To validate this model, we cultured U251 cells using the same patterns. As predicted, F-actins accumulated at the periphery (Fig. S4*I*) and corners of the cells in a corner angle-dependent manner (Fig. 3*G*).

We further examined the effect of inhibiting SpTA via knocking out shootin1b. Interestingly, shootin1b KO not only reduced the translocation velocity (Fig. 1*H*) but also the lifetime (Fig. S4*A*-*C*) of SpTAs. Introduction of these parameters (i.e., SpTA velocity and lifetime derived from shootin1b KO cells) into the model disrupted actin accumulation at the cell periphery and corners (Fig. 3*H* and Movie S10). Consistently, shootin1b KO inhibited actin accumulation at the cell periphery and corners of U251 cells (Fig. 3*I*). Together, these data indicate that SpTAs spontaneously accumulate at cell periphery and protrusions.

### SpTAs drive spontaneous formation of the leading edge for cell polarization

Next, the role of SpTAs in cell morphogenesis was analyzed using U251cells with the deformable plasma membrane. Wild-type (WT) cells typically exhibited a polarized morphology with a large lamellipodium (Fig. 4*A*). On the other hand, inhibition of SpTAs by shootin1b KO resulted in the fragmentation of lamellipodia (Fig. 4*A*), leading to a decrease in the lamellipodial coverage ratio at the cell periphery (Figs. 4*B* and S5*A*). Accordingly, the unilateral integration of lamellipodium was disrupted, resulting in a decrease in the degree of cell polarity (Fig. S5*B*). To analyze how SpTAs drive cell polarization, we first depolymerized F-actins in U251 cells using latrunculin A, and then monitored the cells’ recovery after latrunculin A wash out (33). Treatment of WT U251 cells with 100 nM latrunculin A disrupted their polarized morphology (Fig. 4*C* and Movie S11). During recovery following latrunculin A washout, the cells formed multiple foci of small lamellipodia enriched with F-actins (asterisks, Fig. 4*C*). The foci then grew and moved laterally (cyan arrows) or expanded laterally (yellow arrows), integrating into a large lamellipodium. Through these processes, the cells eventually formed a leading edge and acquired polarity.

**Fig. 4.**
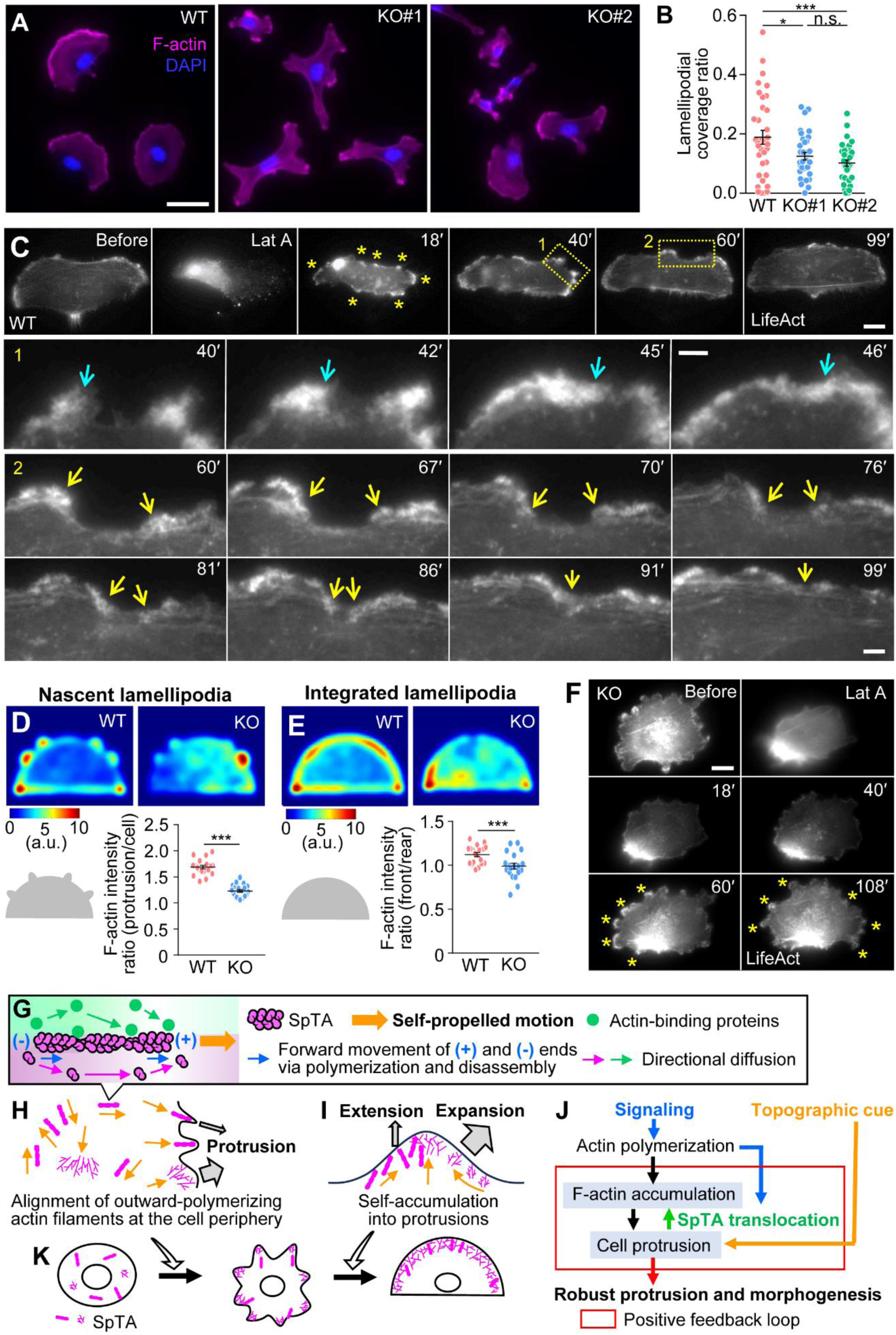
SpTAs drive spontaneous formation of the leading edge for cell polarization. (*A*) WT, shootin1b KO#1 and KO#2 U251 cells stained with Alexa Fluor 594 conjugated phalloidin and DAPI. Scale bar, 50 µm. (*B*) Lamellipodial coverage rate of WT, shootin1b KO#1 and KO#2 U251 cells in (*A*) (WT, KO#1, KO#2, N = 3 experiments, n = 35 cells). Scale bar, 5 µm. Data represent means ± SEM; ***p < 0.01; *p < 0.05; ns, not significant. Statistical analyses were performed using the two-tailed unpaired Student′s *t*-test (KO#1 vs KO#2) and two-tailed unpaired Welch′s *t*-test (WT vs KO#1, WT vs KO#2). (*C*) Fluorescence time-lapse images of a WT U251 cell expressing LifeAct-mCherry obtained by epifluorescence microscopy: lower panels indicate enlarged time-lapse images of the areas indicated by the dashed yellow boxes 1 and 2. The images were taken before and after the treatment with 100 nM latrunculin A (Lat A), and at the indicated times after the Lat A washout. The asterisks indicate actin filament-enriched nascent lamellipodia. Yellow and blue arrows indicate enlargement and lateral expansions of nascent lamellipodia, respectively. See Movie S11. Scale bars, 20 µm; 5 µm (enlarged views). (*D* and *E*) Mathematical model data showing SpTA localization in WT and shootin1b KO cells bearing nascent lamellipodia (*D*) and integrated lamellipodium (*E*), presented in arbitrary unit (a.u.). The panels below describe the cell shape (40 µm width). The graphs below show quantitative data for SpTA accumulation at the nascent lamellipodia (*D*) and integrated lamellipodia (*E*) of WT and shoootin1b KO cells (WT, KO, n = 20 model data). Data represent means ± SEM; ***p < 0.01. Statistical analyses were performed using the two-tailed unpaired Student′s *t*-test. (*F*) Fluorescence time-lapse images of a shootin1b KO#1 U251 cell expressing LifeAct-mCherry obtained by epifluorescence microscopy. The images were taken before and after the treatment with 100 nM Lat A, and at the indicated times after the Lat A washout. The asterisks indicate nascent lamellipodia. See Movie S11. Scale bar, 20 µm. See also Fig. S5*C*-*E*. (*G*) A model of the translocation mechanisms of SpTAs. During the translocation, the front and rear ends of SpTAs move forward via polymerization and disassembly, respectively (blue arrows). Shootin1b facilitates the SpTA translocation by impeding their retrograde flow (Fig. 1F left). As the local concentrations of actin subunits and actin-binding proteins decrease at the front by polymerization and increase at the rear by disassembly, they diffuse along the local concentration gradients (magenta and green arrows). Accompanied by the directional diffusion, SpTAs mediate the translocation of actin and actin-binding proteins. (*H*) SpTA arrival at the cell periphery aligns randomly oriented F-actins towards outward polymerization, which in turn pushes the membrane to protrude (gray arrows). (*I*) SpTAs accumulate at cell protrusions (yellow arrows), leading to their further growth (gray arrows). (*J*) Accumulation of SpTAs at protrusions (green arrow) constitutes a positive feedback interaction with actin-mediated cell protrusion (red box), thereby conferring robustness to the protrusions (red arrow) cell morphogenesis. SpTAs are regulated under cell signaling (blue arrows). This system can also respond to extracellular topographic cues that deform the plasma membrane from outside (yellow arrow). (*K*) Processes of spontaneous polarization of glioma cells. Through local protrusion (*H*) and lateral expansion (*I*), nascent lamellipodia are formed and integrated into a large leading edge for polarization.

We further monitored the motion of SpTAs in a two-dimensional (2D) model of cells bearing protrusions that mimic the nascent and integrated lamellipodia. SpTAs in the model highly accumulated in the nascent lamellipodia (Figs. 4*D* and S5*C*) and moderately accumulated in the integrated lamellipodium located at the convex leading edge (Fig. 4*E* and S5*D*). Introduction of the parameters derived from shootin1b KO cells (Figs. 1*H* and S4*A*-*C*) into the model inhibited SpTA accumulation at the nascent and integrated lamellipodia (Fig. 4*D* and *E*).

Consistently with the modeling data, shootin1b deletion in glioma cells inhibited the lateral movement, expansion, and integration of nascent lamellipodium (asterisks, Fig. 4*F* and Movie S11). This in turn delayed the onset of cell migration after recovery from the latrunculin A treatment (Fig. S5*E*). As previously reported, shootin1b also mediates the generation of traction force at the leading edge through its interaction with treadmilling F-actins (28). Therefore, the impairment of lamellipodial integration resulting from shootin1b KO can be attributed to the combined effects of reduced accumulation of SpTAs at lamellipodia (Fig. 4*D* and *E*) and diminished force generation there. Taken together, we conclude that SpTAs of glioma cells drive spontaneous formation of the leading edge by expanding the nascent lamellipodia.

## DISCUSSION

Treadmilling is an inherent property of actin dynamics. The present study reports SpTAs that translocate within cells as self-propelled “particles” rather than “wave”-like propagation of chemical reactions. The arrival of SpTAs at the cell periphery pushes the membrane to form protrusions. Furthermore, SpTAs accumulate into protrusions, leading to their robust growth and spontaneous formation of cell polarity for migration.

### SpTA translocation by treadmilling

During SpTA translocation (yellow arrow, Fig. 4*G*), their front move forward (blue arrow) via incorporation of ATP-bound actin subunits at the plus end. This is followed by ATP hydrolysis within the filaments, which in turn leads to disassembly of the filaments from the minus ends and forward movement of the rear (blue arrow). The ADP bound to the disassembled actin subunits is replaced with ATP, allowing the subunits to be reused in polymerization (magenta arrows). Shootin1b facilitates SpTA translocation by impeding their retrograde flow (Fig. 1*F* left). During SpTA translocation, local concentrations of actin subunits and actin-binding proteins decrease at the front by polymerization and increase at the rear by disassembly, resulting in their diffusion along the local concentration gradients (green and magenta arrows, Fig. 4*G*). Accompanied by this directional diffusion, SpTAs translocate actin and actin-binding proteins.

SpTAs potentially explain a wide range of intracellular actin dynamics. First, dynamic actin filaments undergo wave-like propagation in various cells (10–13). Their propagations have been mainly explained by the reaction-diffusion model or Turing model, where the excitable component that stimulates actin polymerization at the front simultaneously stimulates the inhibitory component at the rear to depolymerize F-actins (11–13, 34). A variety of signaling and mechanical networks have been proposed to link the excitable and inhibitory components, but a general mechanism has not yet been established (13). Since ATP hydrolysis is an intrinsic property of the actin molecules incorporated in the filaments, stimulation of actin polymerization in SpTAs spontaneously leads to the F-actin disassembly at the rear. By directly linking the actin polymerization at the front to the depolymerization at the rear, treadmilling could be a versatile mechanism for F-actin assembly translocation.

Second, translocation of F-actin assemblies to the cell periphery has been reported to induce the lamellipodia formation (35, 36), as SpTAs did here (Fig. 2*A-C*). Since the polymerizing ends of SpTAs are located at the front (Fig. 4*G*), the arrival of SpTAs at the cell periphery aligns the polymerizing ends outward, thereby pushing the membrane to form filopodia and lamellipodia. Thus, SpTAs explains how the arrival of F-actin assemblies forms filopodia and lamellipodia (Fig. 4*H*).

Third, the accumulation of SpTAs in protrusive regions (Fig. 3*C*-*G*) is a key property of self-propelled particles colliding with a boundary (14). When SpTAs encounter the membrane border, they further undergo lateral translocation, depending on the direction of F-actins relative to the membrane. This lateral movement traps outward-polymerizing F-actins and actin binding proteins at protrusive regions (yellow arrows, Fig. 4*I*) and explains previous observations that different cell types confined to square adhesive islands preferentially extend filopodia and lamellipodia from their protrusive areas (37). This mechanism is distinct from the previously reported actin accumulation mechanism via curvature-sensitive actin nucleation (38, 39).

### SpTAs in spontaneous cell protrusion and morphogenesis

The F-actin accumulation at cell protrusions through SpTA translocation (green arrow, Fig. 4*J*) constitutes a positive feedback interaction with the F-actin-driven cell protrusion (red box). This interaction thereby confers robustness to the protrusions (red arrow) and leads to their further growth (gray arrows, Fig. 4*I*). Since translocation of SpTAs into the protrusion (yellow arrows, Fig. 4*I*) results in their lateral decrease, the self-propelled motion of SpTAs also suppresses lateral protrusive activities. During spontaneous polarization of glioma cells, nascent protrusions were integrated into a large leading edge through their lateral movement and expansion (Fig. 4*K*), accompanied by a relative decrease in SpTAs at the rear (Fig. 4*C*, Movie S11). These processes would work in concert with other cytoskeletal dynamics, including actomyosin contraction at the rear (23, 40) to spontaneously polarize cells and initiate cell migration. Spontaneous polarization would also be supported by membrane tension generated by actin-based protrusions (41, 42) as well as by membrane deformation through curved membrane-bound proteins with adhesion (39).

F-actins undergo outward polymerization not only at the filopodia and lamellipodia but also in various cell protrusions, such as growth cones and dendritic spines of neurons (43, 44), microvilli (4, 5), cell-cell adhesion sites (45) and microridge (46) of epithelial cells, hair cell stereocilia (47), cancer cell invadopodia (3), and phagocytic cups (3). Our modeling data showed that SpTAs self-accumulate at small protrusive and large convex regions (Fig. 4*D* and *E*), as well as protrusions with different apex angles (Fig. 3*F*). Different actin-binding proteins and physical interactions could control the mode of protrusion, either extension or expansion, by altering the shape of the F-actin assembly (2, 3, 48, 49). Thus, SpTAs have the potential to form a wide range of actin-based cell protrusions.

### SpTAs in environmental cue-driven cell morphogenesis

Because actin polymerization and disassembly are regulated by signaling pathways (1–3, 6, 8, 9), the formation, translocation, and disappearance of SpTAs are subject to the regulation by extracellular chemical cues and cell signaling (blue arrows, Fig. 4*J*). Inhibition of the Arp2/3 complex or formins reduced the SpTA translocation velocity (Fig. 1*G*), indicating that SpTAs are under the regulation of these molecules. The Arp2/3 complex and formins are located at the plasma membrane, and a recent study reported that formins attached to intracellular vesicles promote actin polymerization in the cytoplasm (50). These actin regulators would maintain the polymerizing ends of SpTAs along the plasma membrane and in the cytoplasm. Additionally, SpTAs could mediate cellular responses to extracellular topographic cues that deform the plasma membrane from outside of the cell (yellow arrow, Fig. 4*J*), as they accumulate at the protrusive and convex regions. This allows migrating cells to navigate flexibly through mechanical barriers and tissue pores in three-dimensional environments (34, 51, 52). The roles of SpTAs in various actin-based cell protrusions, as well as their regulation by environmental cues that tune spontaneous cell morphogenesis, are important issues for future analyses.

## Materials and Methods

### Cell culture and transfection

A subline of glioma U251 cells (23) and COS7 cells were seeded on glass-bottom dishes (Matsunami, #D11130H) coated with 100 μg/ml poly-D-lysine (Sigma) and 5 μg/ml laminin (Lonza), and cultured in DMEM high glucose medium (Nacalai, #08458-16) supplemented with 10% fetal bovine serum (Japan Bio Serum) at 37°C and 5% CO_2_. They were transfected with vectors using Polyethylenimine MAX (PEI MAX, Polysciences) following the manufacturer’s protocol.

### Generation of shootin1b KO U251 cells by CRISPR-Cas9

The guide oligonucleotide sequence (forward: 5’-CACCGGCAGCTCATTACCAGTCTGA-3’; reverse: 5’-AAACTCAGACTGGTAATGAGCTGCC-3’), corresponding to nucleotides 30-49 in the coding region of human shootin1b, were selected by an online CRISPR Cas9 Design tool (http://crispr.mit.edu/). The pSpCas9 (BB)-2A-Puro (PX459) V2.0 plasmid (Addgene, #62988) was linearized using by BbsI-HF (New England Biolabs, #R3539S), then the guide oligonucleotides were annealed and inserted into the vector using Ligation high Ver.2 (TOYOBO, #LGK-201). U251 cells were transfected with the plasmid, cultured individually in 96-well plates, and screened using 4 μg/ml puromycin. Shootin1b depletion in KO cells was confirmed by immunoblot analyses (Fig. S2*G*).

### Micropatterning of laminin on the substrate for cell adhesion

Micropatterns of laminin for cell adhesion were produced on a glass surface following our previously established procedure (53), with modifications (Fig. S3). The glass-bottom dish (Iwaki, #3961-035) was cleaned using ultrasonic waves for 5 min with special grade ethanol, followed by rinsing with water and additional 5-min ultrasonic cleaning with pure water. After the cleaning, the dish was air-dried in a clean room at room temperature. 3-Phenylaminopropyltrimethoxysilane (2%, Shin-Etsu, #LS-4500) was dissolved in a 2% solution of acetic acid in water and then poured onto the glass-bottom dish, followed by incubation for 30 min. After being washed twice with water, the dish was dried at 70°C for 1 h. This process resulted in the covalent bonding of the 3-phenylaminopropyl group with the glass surface, which made it hydrophobic. Any residues resulting from the reaction were eliminated by ultrasonically cleaning with special grade ethanol for 5 min.

The hydrophobic glass surface was coated with 100 µL of ethanol containing 0.2% of MPC (2-methacryloyloxyethyl phosphorylcholine) polymer (NOF, #Lipiduro-CM5206). After the reaction for 10-15 sec, the coating solution was removed and left to dry overnight at room temperature in the dark. This resulted in the formation of an MPC polymer film on the glass (Fig. S3). Then, the dish was incubated with 2 ml H_2_O overnight. The cell non-adhesive MPC-polymer layer was removed by a programmed raster scanning of a femtosecond laser (center wavelength 800 nm, pulse width 130 fsec, pulse energy 100 nJ, transmission frequency 5 kHz, Solstice, Spectra-Physics) focused with an objective lens 20x 0.46 NA (Olympus, UMPlanFl) at a speed of 400 µm/s on the glass surface. The raster scanning patterns are shown (Figs. S3 and S4F). Then, the cell-adhesive patterns of laminin were produced by coating the exposed glass surface with poly-D-lysine (PDL) and laminin. The cell-adhesive patterns were confirmed by coating with green fluorescent-HiLyte 488-Laminin (Cytoskeleton, #LMN02-A) (Fig. S3).

### Immunocytochemistry

Immunocytochemistry was performed as described previously (25). Cultured cells were fixed with 3.7% formaldehyde in Krebs buffer (118 mM NaCl, 4.7 mM KCl, 1.2 mM KH_2_PO_4_, 1.2 mM MgSO_4_, 4.2 mM NaHCO_3_, 2 mM CaCl_2_, 10 mM glucose, 400 mM sucrose, 10 mM HEPES pH 7.0) for 20 min on ice and 10 min at room temperature, followed by treatment for 15 min with 0.05% Triton X-100 in PBS on ice and 10% fetal bovine serum in PBS for 1 h at room temperature. The cells were incubated with a primary antibody diluted in PBS containing 10% fetal bovine serum overnight at 4°C. They were washed with PBS, and then incubated with secondary antibody diluted in PBS for 1 h at room temperature. After washing with PBS, the cells were stained with Alexa Fluor 594 conjugated phalloidin (1:100, Thermo Fisher Scientific) and DAPI (1:1000, Thermo Fisher Scientific) for 30 min at room temperature. The fluorescence and phase-contrast images of U251 cells were acquired a fluorescence microscope (Carl Zeiss, Axioplan2) equipped with an objective lens 40x 0.75 NA (Carl Zeiss, plan-Neofluar), a charge-coupled device camera (Carl Zeiss, AxioCam MRm) and imaging software (Carl Zeiss, ZEN), and a TIRF microscope (Olympus, IX81) equipped with a CMOS camera (Hamamatsu, ORCA Flash4.0LT), an objective lens 100x 1.49 NA (Olympus, UAPON), and MetaMorph software.

### Immunoblot

Cultured U251 cells were lysed with RIPA buffer (50 mM Tris-HCl, pH 8.0), 1 mM EDTA, 150 mM NaCl, 1% Triton X-100, 0.1% SDS, 0.1% sodium deoxycholate, 1 mM DTT, 1 mM PMSF, and 0.01 mM leupeptin), and incubated for 10 min at 4°C. Subsequently, the cell lysate was centrifuged at 15000 rpm for 15 min at 4°C. The supernatant was mixed with an equal volume of 2 x SDS sample buffer (131 mM Tris-HCl, pH 6.8, 21% glycerol, 4% SDS, 0.05% bromophenol blue and 5% β-mercaptoethanol). The mixture was incubated for 5 min at 95°C, followed by SDS-polyacrylamide gel electrophoresis. Immunoblotting was performed as described previously (28).

### Antibodies

The following primary antibodies were used: anti-shootin1 (rabbit, 1:400) (54); anti-shootin1b (rabbit, 1:2000) (54); anti-NCAM-L1 (mouse, 1:1000, Santa Cruz Biotechnology, #sc-514360); anti-cortactin (mouse, 1:1000, Millipore, #05-180); anti-actin (mouse, 1:1000, Millipore, #MAB1501R); anti-ARPC2 (rabbit, 1:1000, Millipore, #07-227). The following secondary antibodies were used: anti-rabbit (Alexa 594 conjugated donkey, 1:1000, Jackson ImmunoResearch Labs, #711-585-152); anti-rabbit (Alexa 488 conjugated goat, 1:1000, Thermo Fisher Scientific, #A-11008); anti-mouse (Alexa 488 conjugated goat, 1:1000, Thermo Fisher Scientific, #A-11029); anti-rabbit (HRP conjugated donkey, 1:2000, GE Healthcare, #NA934); anti-mouse (HRP conjugated goat, 1:5000, Thermo Fisher Scientific, #AP308P).

### DNA construction

To generate pEGFP-N1-LifeAct vector, the annealed LifeAct (forward: 5’-TCGAGATGGGTGTCGCAGATTTGATCAAGAAATTCGAAAGCATCTCAAAGGAAGAAGGG-3’; reverse: 5’-GATCCCTTCTTCCTTTGAGATGCTTTCGAATTTCTTGATCAAATCTGCGACACCCATC-3’) was fused to N-terminal of EGFP-N1 vectors (Clontech). To generate pLifeAct-mCherry-N1 vector, the annealed LifeAct was fused to N-terminal of mCherry-N1 vectors (Takara, #632523). Generation of pFN21A-HaloTag-actin was described previously (28).

### TIRF and epifluorescence microscopy

U251 cells were cultured on glass-bottom dishes coated with PDL and laminin. Actin dynamics in close proximity to the ventral plasma membrane were visualized using the F-actin marker EGFP-LifeAct or LifeAct-mCherry under TIRF or epifluorescence microscopy. In addition, to differentially visualize SpTAs translocating in close proximity to the ventral plasma membrane and those detached from the membrane, we combined TIRF microscopy, which visualizes the region in close proximity to the ventral plasma membrane, and epifluorescence microscopy, which visualizes the entire cell thickness (Fig. S1*E*). U251 cells expressing LifeAct-mCherry and EGFP-LifeAct were cultured on glass-bottom dishes coated with PDL and laminin. Prior to observation, the cells were incubated in DMEM/F-12 Ham medium (Sigma) supplemented with 10% FBS (Japan Bio Serum). The TIRF laser was used to observe fluorescence images of LifeAct-mCherry, while the epifluorescence laser was used to observe fluorescence images of EGFP-LifeAct. The observations were made with a TIRF microscope (Olympus, IX81) equipped with a CMOS camera (Hamamatsu, ORCA Flash4.0LT), an objective lens 100x 1.49 NA or 150x 1.45 NA (Olympus, UAPON), and MetaMorph software. Images were acquired every 10 or 60 sec.

### Fluorescent speckle imaging of SpTA by TERF microscopy

Actin dynamics within SpTAs were examined the by labelling F-actins with EGFP-LifeAct, and monitoring actin molecules in SpTAs by speckle imaging of HaloTag-actin (24). U251 cells expressing HaloTag-actin and EGFP-LifeAct were cultured on glass-bottom dishes coated with PDL and laminin. Prior to observation, the cells were treated with 0.05 μM HaloTag TMR ligand (Promega) in the culture medium and incubated for 1 h at 37°C. The ligand was washed with PBS, and the cells were then incubated in DMEM/F-12 Ham medium (Sigma, #D8062) supplemented with 10% FBS (Japan Bio Serum). The fluorescent speckles of HaloTag-actin and EGFP-LifeAct were observed using a TIRF microscope (Olympus, IX81) equipped with a CMOS camera (Hamamatsu, ORCA Flash4.0LT), an objective lens 150x 1.45 NA (Olympus, UAPON), and MetaMorph software. Fluorescence images were acquired every 10 sec. Actin flow velocity was calculated by tracing the speckles of HaloTag-actin in SpTAs using ImageJ (Fiji version). Actin polymerization rate was calculated as the sum of SpTA migration velocity, visualized by EGFP-LifeAct, and HaloTag-actin flow velocity, as reported (24).

### 3D imaging by confocal deconvolution microscopy

U251 cells expressing HaloTag-actin and EGFP-LifeAct were cultured in 2D (on glass-bottom dishes coated with PDL and laminin) or 3D (75% matrigel) environment. Before observation, the cells were incubated with 0.05 μM HaloTag TMR ligand (Promega) in the culture medium and incubated for 1 h at 37°C. The ligand was washed with PBS, and the cells were then incubated in DMEM/F-12 Ham medium (Sigma) supplemented with 10% FBS (Japan Bio Serum). The fluorescent speckles of HaloTag-actin and EGFP-LifeAct were observed using a confocal microscope (Leica, Stellaris 8) equipped with an objective lens 100x 1.40 NA (Leica, HC PL APO CS2) and Las X software. Deconvolution of 3D image stacks were obtained by Las X software.

### Mathematical modelling

To investigate whether SpTAs mediate the accumulation of F-actins at the protrusions of cells confined to adhesive islands (Fig. 3*C*), we constructed a mathematical model that describes the movement of intracellular SpTAs. Our model is based on quantitative data obtained from experimental observations: we focused on filopodium-type SpTAs because it is difficult to observe individual F-actins in lamellipodium-type SpTAs, and filopodium-type and lamellipodium-type SpTAs are interchangeable (Fig. 1*B*). For simplicity, the present model describes the translocation of the SpTAs, not a detailed mechanism of SpTA translocation (25). The model depicts the processes of SpTA emergence and disappearance, as well as changes in the direction of SpTA translocation.

### Emergence and disappearance of SpTAs

Our experimental data indicate that SpTAs emerge widely in the cytoplasm, move in random directions, and then disappear (Movie S1). In our model, the emergence locations and translocation directions of SpTAs are randomly chosen at the start of the modeling. The model data are updated every 1 sec (Δt = 1 sec), generating one SpTA in the cell with a probability of 10%. The lifetimes of SpTAs in WT and shootin1b KO cells, quantified by live cell imaging, were modeled using gamma distributions (Fig. S4*A* and *B*).

The length (*L*) and translocation velocity (*V*) of SpTAs are also derived from experimental data (Fig. 1*H* and Fig. S4*E*); model uses their average values for simplicity. SpTAs located at the cell periphery move laterally along the cell membrane (Fig. 2*A*). The velocity of the lateral movement was modelled as cos*θ* of the mean velocity of SpTAs (Fig. 1*H*), where *θ* is the angle of the direction of SpTA movement with respect to the cell membrane (Fig. 2*D*).

The local concentration of actin subunits increases immediately after the disappearance (complete disassembly) of SpTA (Fig. 2*C*). Since actin polymerization depends on the local concentration of actin subunits, our model assumes that an SpTA reemerges at the location where the preceding SpTA disappeared and translocates in a random direction.

### Change in the direction of SpTA translocation

SpTAs frequently change the direction of their translocation (Movies S1). The change in direction was quantified every 50 sec from live cell imaging data (Fig. S4*D*). As the angular distribution of these changes was symmetric with a zero peak, we modelled it using a Gaussian distribution (Fig. S4*D*).

### Model parameters estimated from experimental data

- Translocation velocity of SpTA (Fig. 1*H*): 2.0 µm/min (WT), 1.5 µm/min (KO#1)
- SpTA length (Fig. S4*E*): 4.1 µm
- Parameters of the gamma distribution representing lifetime variation; estimated from quantitative data (Fig. S4*A* and *B*): α = 3.0, β = 119 (WT), α = 2.4, β = 106 (KO#1)
- Parameters of the Gaussian distribution representing SpTA directional change; quantified from experimental data (Fig. S4*D*): μ = −0.61, σ = 11

### Analyses of F-actin accumulation and lamellipodial formation

Accumulation of F-actins at the cell periphery was determined by dividing the average fluorescence intensity of F-actins within a 2 μm-wide region from the cell periphery (green region, Fig. S4*G*) by the average intensity of F-actins in the whole cell region (blue region). Accumulation of F-actins at the cell protrusions was determined by dividing the average intensity of F-actins at the corner regions (yellow region: 2 μm-wide and 5 μm-long regions in both sides of the corner, Fig. S4*G*) by the average intensity of F-actins in the peripheral region (green region). Accumulation of F-actins at the nascent lamellipodia was determined by dividing the average fluorescence intensity of F-actins in protrusive regions (green regions, Fig. S5*C*) by the average intensity of F-actins in the whole cell region (blue region). Accumulation of F-actins at the integrated lamellipodium was determined by dividing the average intensity of F-actins at the convex leading edge (yellow region: 4 μm-wide, Fig. S5*D*) by the average intensity of F-actins in the rear region (green region: 4 μm-wide). Lamellipodial coverage rate was calculated by dividing the total length of lamellipodia (yellow line, Fig. S5*A*) by the perimeter of the cell (green line). The fluorescence intensity of F-actins was analyzed using ImageJ (Fiji version). Lamellipodia was defined as the area of cell periphery where the fluorescence intensity exceeded the mean intensity + SD of the whole cell body. The polar histogram of the lamellipodial areas was generated using MATLAB (Fig. S5*B*).

### Quantification and statistical analyses

Statistical analyses were performed using Excel (Microsoft 365) or GraphPad Prism 7 (GraphPad Software). For samples with more than 7 data points, the D′Agostino–Pearson normality test was used to determine whether the data followed a normal distribution. In cases where the number of data points was between 3 and 7, the Shapiro‒Wilk test was used for the normality test. We also tested the equality of variation with the F test for two independent groups that followed normal distributions. Significance tests were performed as follows: (1) two-tailed unpaired Student′s *t*-test to compare normally distributed data with equal variance from two independent groups; (2) two-tailed unpaired Welch′s *t*-test to compare normally distributed data with unequal variance from two independent groups; (3) two-tailed Mann–Whitney *U*-test to compare nonnormally distributed data from two independent groups. The statistical information and number of samples for each experiment are indicated in the Fig. legends. For detailed statistical results including the test statistics and exact p values, see the statistical source data associated with each Figure. All data are shown as the mean ± SEM. Statistical significance was defined as ***p < 0.01; **p < 0.02; *p < 0.05; ns, not significant. All experiments were performed at least three times and reliably reproduced. The investigators were blinded to the experimental groups for each analysis.

## Data, Materials and Code Availability

All study data are included in the article and/or Supplementary Information. Source data of immunoblot and statistical analyses in the Figs. have been deposited at Mendeley Data (DOI: 10.17632/ymmcbh92v6.1) and are publicly available as of the date of publication. The MATLAB codes used for mathematical modeling are available at GitHub (https://github.com/KioYagami/actin-wave/tree/master). Any additional information required to reanalyze the data reported in this paper is available upon reasonable request.

## Acknowledgments

We thank Drs. Masatoshi Takeichi (Riken) and Shintaro Suzuki (Kwansei Gakuin Univ) for providing the subline of U251 cells, Drs. Hitoshi Sakano (Fukui Univ) and Nir Gov (Weizmann Inst Sci) for reading the manuscript, Drs. Jim Bamburg (CSU), Robert Grosse (University of Freiburg), Carsten Beta (Potsdam Univ), Sean Collins (UC Davis), Klemens Rottner (Tech Univ Braunschweig), Giovanni Volpe (Univ Gothenburg), Shiro Suetsugu (NAIST), Naoki Watanabe (Kyoto Univ) and Yusuke Maeda (Kyushu Univ) for valuable discussions, and Satoko Shimamura for kind encouragements. This research was supported in part by AMED under Grant Number JP17gm0810011 (N.I.), JSPS KAKENHI (JP19H03223, N.I.), JSPS Grants-in-Aid for Early-Career Scientists (JP19K16127, K.B.), and the Osaka Medical Research Foundation for Incurable Diseases (K.B., T.M.).

## References

1. T. D. Pollard, G. G. Borisy, Cellular motility driven by assembly and disassembly of actin filaments. Cell 112, 453–465 (2003).

2. M. F. Carlier, S. Shekhar, Global treadmilling coordinates actin turnover and controls the size of actin networks. Nat. Rev. Mol. Cell Biol. 18, 389–401 (2017).

3. K. Rottner, M. Schaks, Assembling actin filaments for protrusion. Curr. Opin. Cell Biol. 56, 53–63 (2019).

4. N. S. Gov, Dynamics and morphology of microvilli driven by actin polymerization. Phys. Rev. Lett. 97, 018101 (2006).

5. L. M. Meenderink et al., Actin dynamics drive microvillar motility and clustering during brush border assembly. Dev. Cell 50, 545–556 e544 (2019).

6. B. W. Bernstein, J. R. Bamburg, ADF/cofilin: a functional node in cell biology. Trends Cell Biol. 20, 187–195 (2010).

7. E. A. Vitriol, A. L. Wise, M. E. Berginski, J. R. Bamburg, J. Q. Zheng, Instantaneous inactivation of cofilin reveals its function of F-actin disassembly in lamellipodia. Mol. Biol. Cell 24, 2238–2247 (2013).

8. Y. Wang et al., Formin-like 2 Promotes beta1-Integrin Trafficking and Invasive Motility Downstream of PKCalpha. Dev .Cell 34, 475–483 (2015).

9. B. L. Goode, J. Eskin, S. Shekhar, Mechanisms of actin disassembly and turnover. J. Cell Biol. 222 (2023).

10. A. E. Carlsson, Actin dynamics: from nanoscale to microscale. Annu. Rev. Biophys. 39, 91–110 (2010).

11. J. Allard, A. Mogilner, Traveling waves in actin dynamics and cell motility. Curr. Opin. Cell Biol. 25, 107–115 (2013).

12. N. Inagaki, H. Katsuno, Actin waves: origin of cell polarization and migration? Trends Cell Biol. 27, 515–526 (2017).

13. C. Beta, L. Edelstein-Keshet, N. Gov, A. Yochelis, From actin waves to mechanism and back: How theory aids biological understanding. eLife 12, e87181 (2023).

14. A. Kaiser, H. H. Wensink, H. Lowen, How to capture active particles. Phys. Rev. Lett. 108, 268307 (2012).

15. J. Elgeti, R. G. Winkler, G. Gompper, Physics of microswimmers--single particle motion and collective behavior: a review. Rep. Prog. Phys. 78, 056601 (2015).

16. C. Bechinger et al., Active particles in complex and crowded environments. Rev. Mod. Phys. 88, 045006 (2016).

17. S. J. Ebbens, J. R. Howse, In pursuit of propulsion at the nanoscale. Soft Matter 6, 726–738 (2010).

18. M. Paoluzzi, R. Di Leonardo, M. C. Marchetti, L. Angelani, Shape and displacement fluctuations in soft vesicles filled by active particles. Sci. Rep. 6, 34146 (2016).

19. Y. Li, P. R. Ten Wolde, Shape transformations of vesicles induced by swim pressure. Phys. Rev. Lett. 123, 148003 (2019).

20. H. R. Vutukuri et al., Active particles induce large shape deformations in giant lipid vesicles. Nature 586, 52–56 (2020).

21. J. F. Boudet et al., From collections of independent, mindless robots to flexible, mobile, and directional superstructures. *Sci*. Robot. 6, eabd0272 (2021).

22. W. Oosterheert, B. U. Klink, A. Belyy, S. Pospich, S. Raunser, Structural basis of actin filament assembly and aging. Nature 611, 374–379 (2022).

23. V. Vassilev, A. Platek, S. Hiver, H. Enomoto, M. Takeichi, Catenins steer cell migration via stabilization of front-rear polarity. Dev. Cell 43, 463–479 (2017).

24. T. Minegishi, R. Fujikawa, R. F. Kastian, Y. Sakumura, N. Inagaki, Analyses of actin dynamics, clutch coupling and traction force for growth cone advance. J. Vis. Exp., e63227 (2021).

25. H. Katsuno et al., Actin migration driven by directional assembly and disassembly of membrane-anchored actin filaments. Cell Rep. 12, 648–660 (2015).

26. Y. Kubo et al., Shootin1-cortactin interaction mediates signal-force transduction for axon outgrowth. J. Cell Biol. 210, 663–676 (2015).

27. K. C. Flynn, C. W. Pak, A. E. Shaw, F. Bradke, J. R. Bamburg, Growth cone-like waves transport actin and promote axonogenesis and neurite branching. Dev. Neurobiol. 69, 761–779 (2009).

28. T. Minegishi et al., Shootin1b mediates a mechanical clutch to produce force for neuronal migration. Cell Rep. 25, 624–639.e626 (2018).

29. B. J. Nolen et al., Characterization of two classes of small molecule inhibitors of Arp2/3 complex. Nature 460, 1031–1034 (2009).

30. Y. Nishimura et al., The formin inhibitor SMIFH2 inhibits members of the myosin superfamily. J. Cell Sci. 134, jcs253708 (2021).

31. L. A. Cameron, J. R. Robbins, M. J. Footer, J. A. Theriot, Biophysical parameters influence actin-based movement, trajectory, and initiation in a cell-free system. Mol. Biol. Cell 15, 2312–2323 (2004).

32. T. Kalwarczyk et al., Comparative analysis of viscosity of complex liquids and cytoplasm of mammalian cells at the nanoscale. Nano Lett. 11, 2157–2163 (2011).

33. G. Gerisch et al., Mobile actin clusters and traveling waves in cells recovering from actin depolymerization. Biophys. J. 87, 3493–3503 (2004).

34. O. D. Weiner, W. A. Marganski, L. F. Wu, S. J. Altschuler, M. W. Kirschner, An actin-based wave generator organizes cell motility. PLoS Biol. 5, e221 (2007).

35. T. Bretschneider et al., The three-dimensional dynamics of actin waves, a model of cytoskeletal self-organization. Biophys. J. 96, 2888–2900 (2009).

36. K. Zhang, W. Lyu, J. Yu, A. J. Koleske, Abl2 is recruited to ventral actin waves through cytoskeletal interactions to promote lamellipodium extension. Mol. Biol. Cell 29, 2863–2873 (2018).

37. K. K. Parker et al., Directional control of lamellipodia extension by constraining cell shape and orienting cell tractional forces. FASEB J. 16, 1195–1204 (2002).

38. J. Saarikangas et al., MIM-Induced Membrane Bending Promotes Dendritic Spine Initiation. Dev. Cell 33, 644–659 (2015).

39. R. K. Sadhu, S. Penič, A. Iglič, N. S. Gov, Modelling cellular spreading and emergence of motility in the presence of curved membrane proteins and active cytoskeleton forces. Eur. Phys. J. Plus 136, 495 (2021).

40. A. Hadjitheodorou et al., Directional reorientation of migrating neutrophils is limited by suppression of receptor input signaling at the cell rear through myosin II activity. Nat. Commun. 12, 6619 (2021).

41. H. De Belly et al., Cell protrusions and contractions generate long-range membrane tension propagation. Cell 186, 3049–3061 e3015 (2023).

42. K. Tsujita, T. Takenawa, T. Itoh, Feedback regulation between plasma membrane tension and membrane-bending proteins organizes cell polarity during leading edge formation. Nat. Cell Biol. 17, 749–758 (2015).

43. D. M. Suter, P. Forscher, Substrate-cytoskeletal coupling as a mechanism for the regulation of growth cone motility and guidance. J. Neurobiol. 44, 97–113 (2000).

44. N. A. Frost, H. Shroff, H. Kong, E. Betzig, T. A. Blanpied, Single-molecule discrimination of discrete perisynaptic and distributed sites of actin filament assembly within dendritic spines. Neuron 67, 86–99 (2010).

45. N. Efimova, T. M. Svitkina, Branched actin networks push against each other at adherens junctions to maintain cell-cell adhesion. J. Cell Biol. 217, 1827–1845 (2018).

46. P. Y. Lam, S. Mangos, J. M. Green, J. Reiser, A. Huttenlocher, In vivo imaging and characterization of actin microridges. PloS one 10, e0115639 (2015).

47. A. K. Rzadzinska, M. E. Schneider, C. Davies, G. P. Riordan, B. Kachar, An actin molecular treadmill and myosins maintain stereocilia functional architecture and self-renewal. J. Cell Biol. 164, 887–897 (2004).

48. L. Huber, R. Suzuki, T. Kruger, E. Frey, A. R. Bausch, Emergence of coexisting ordered states in active matter systems. Science 361, 255–258 (2018).

49. K. F. Swaney, R. Li, Function and regulation of the Arp2/3 complex during cell migration in diverse environments. Curr. Opin. Cell Biol. 42, 63–72 (2016).

50. D. Frank et al., Vesicle-Associated Actin Assembly by Formins Promotes TGFbeta-Induced ANGPTL4 Trafficking, Secretion and Cell Invasion. Adv. Sci. 10, e2204896 (2023).

51. K. M. Yamada, M. Sixt, Mechanisms of 3D cell migration. Nat. Rev. Mol. Cell Biol. 20, 738–752 (2019).

52. A. Hadjitheodorou et al., Leading edge competition promotes context-dependent responses to receptor inputs to resolve directional dilemmas in neutrophil migration. Cell Syst. 14, 196–209 e196 (2023).

53. K. Okano et al., In situ laser micropatterning of proteins for dynamically arranging living cells. Lab Chip 13, 4078–4086 (2013).

54. Y. Higashiguchi et al., Identification of a shootin1 isoform expressed in peripheral tissues. Cell Tissue Rsssses. 366, 75–87 (2016).

