## Supplemental Information for "Spontaneous cell morphogenesis via self-propelled actin filaments"

Naoyuki Inagaki

**This PDF file includes:**

Figures S1 to S5

Legends for Movies S1 to S11

**Other supporting materials for this manuscript include the following:**

Movies S1 to S11

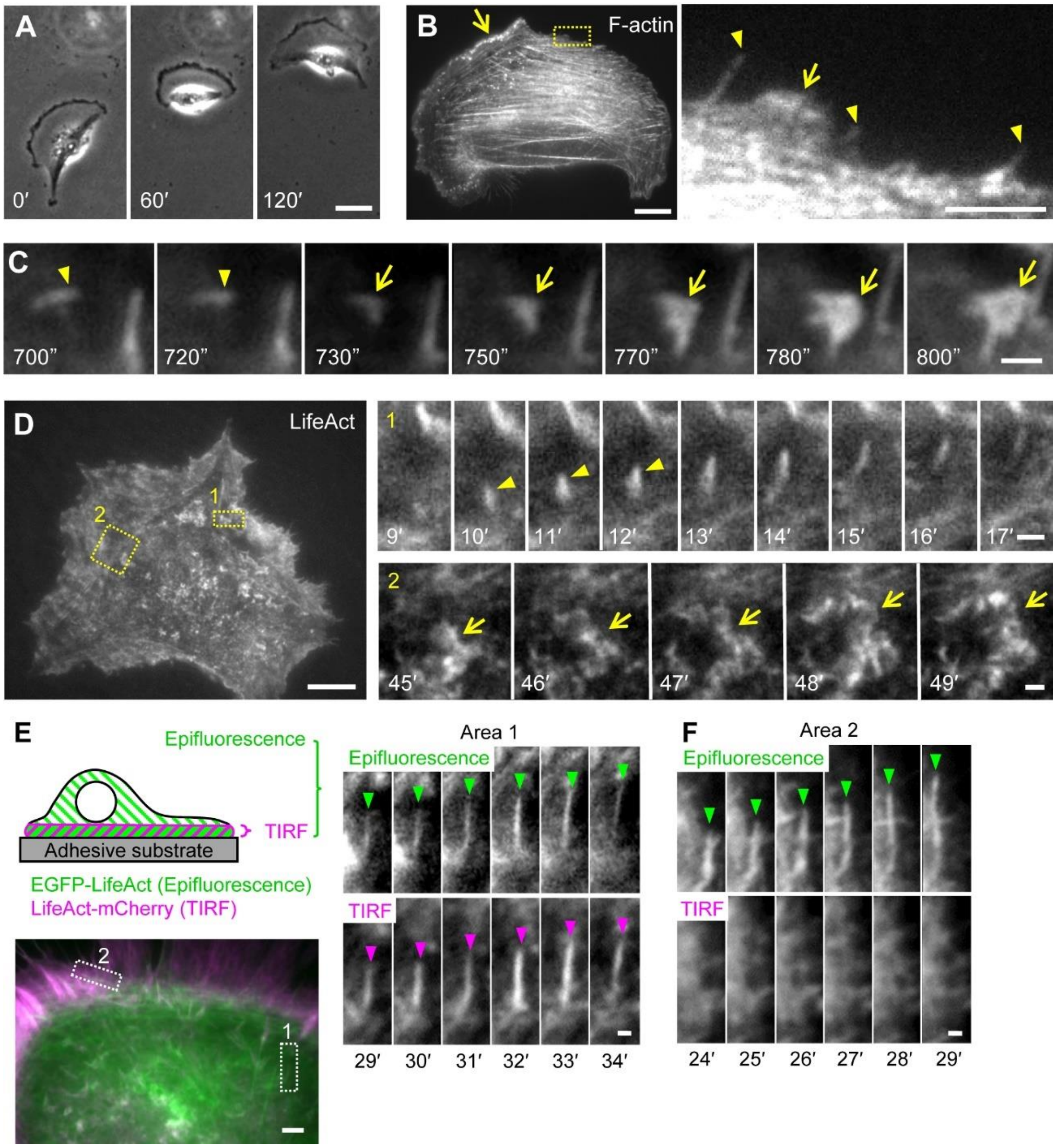


**Fig. S1.** F-actin assemblies translocate widely in cells. (*A*) Time-lapse phase-contrast images of U251 cells. Scale bar, 20 µm. (*B*) A U251 cell stained with Alexa Fluor 594 conjugated phalloidin. An enlarged view of the rectangular region is shown to the right. Arrows and arrowheads indicate lamellipodia and filopodia, respectively. Scale bars, 20 µm (left); 5 µm (right). (*C*) Fluorescence time-lapse images of a lamellipodium-type F-actin assembly (arrows) expanded from a filopodium-type F-actin assembly (arrowheads) in a U251 cell in Fig. 1B. See Movie S1. Scale bar, 2 µm. (*D*) A fluorescence image of a COS7 cell expressing LifeAct-mCherry obtained by TIRF microscopy: right panels indicate enlarged time-lapse images in rectangular regions 1 and 2. Arrows and arrowheads indicate lamellipodium-type and filopodium-type F-actin assemblies, respectively. See Movie S1. Scale bars, 20 µm (left); 2 µm (right). (*E* and *F*) Fluorescence time-lapse images of a U251 cell expressing LifeAct-mCherry (obtained by TIRF microscopy) and EGFP-LifeAct (obtained by epifluorescence microscopy): right panel and (*F*) indicate enlarged time-lapse images in rectangular regions 1 and 2. See Movie S2. Scale bars, 5 µm (*E*); 1 µm (*E* enlarged view, *F*).

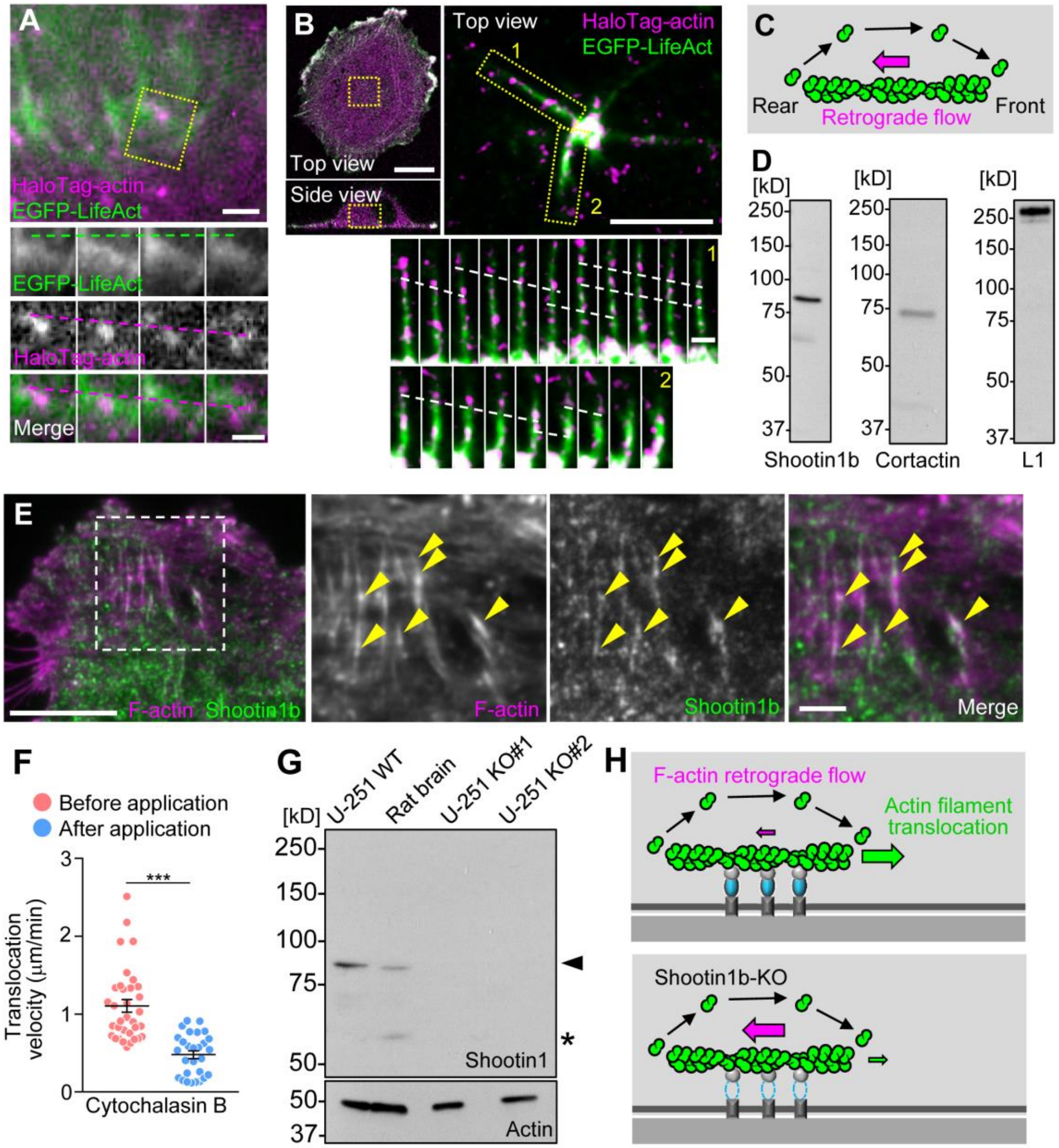
A
HaloTag-actin
EGFP-LifeAct
EGFP-LifeAct
HaloTag-actin
Merge
B
Top view
Side view
Top view
HaloTag-actin
EGFP-LifeAct
1
2
C
Rear
Retrograde flow
Front
D
[kD]
250
150
100
75
50
37
Shootin1b
Cortactin
L1
E
F-actin Shootin1b
F-actin
Shootin1b
Merge
F
Before application
After application
Translocation velocity (μm/min)
3
2
1
0
***
Cytochalasin B
G
U-251 WT
Rat brain
U-251 KO#1
U-251 KO#2
Shootin1
Actin
H
F-actin retrograde flow
Actin filament translocation
Shootin1b-KO

29

**Fig. S2.** (*A*) A fluorescent speckle image of HaloTag-actin in a U251 cell obtained by TIRF microscopy; actin filaments were also monitored by EGFP-LifeAct. Time-lapse montages of a filopodium-type F-actin assembly in the rectangular region at 10-sec intervals are shown below. Green and magenta lines indicate the F-actin front and F-actin retrograde flow, respectively. Scale bar, 1 µm. (*B*) A Fluorescent speckle image of HaloTag-actin in a U251 cell obtained by 3D imaging with confocal deconvolution microscopy; actin filaments were also monitored by EGFP-LifeAct. Dotted lines indicate F-actin retrograde flow. Scale bars, 20 µm (left); 5 µm (upper right); 1 µm (lower right). (*C*) A diagram explaining F-actin retrograde flow accompanied by its polymerization at the front and disassembly at the rear: treadmilling. (*D*) Immunoblot analyses of U251 cells with anti-shootin1b, anti-cortactin and anti-L1 antibodies. (*E*) A fluorescence image of a U251 cell stained with anti-shootin1b antibody and Alexa Fluor 594 conjugated phalloidin (for actin filament) obtained by TIRF microscopy. An enlarged view of the area within the rectangle is shown in the right panel. Arrowheads indicate shootin1b colocalization with actin filaments. Scale bars, 20 µm (left); 5 µm (right). (*F*) Translocation velocities of filopodium-type F-actin assemblies before and after the application of 0.05 µM cytochalasin B (N = 4 cells, before application: n = 34, after application: n = 28). Data represent means ± SEM; ***p < 0.01. Statistical analyses were performed using the two-tailed Mann–Whitney *U*-test. (*G*) Immunoblot analysis of shootin1b in WT U251 cells, postnatal day 1 rat brain, and shootin1b KO U251 cells (KO#1 and KO#2). The cell and tissue lysates were immunoblotted with anti-shootin1 and anti-actin antibodies. The arrowhead and asterisk indicate the bands corresponding to shootin1b and shootin1a, respectively. Shootin1a is a splicing variant of shootin1b expressed in neurons. Immunoblot with anti-actin antibody served as a loading control. (*H*) A diagram explaining the effects of shootin1b KO on the velocities of actin filament retrograde flow and F-actin assembly translocation. If actin filaments in F-actin assemblies are anchored to the plasma membrane and adhesive substrate through shootin1a, this anchoring impedes the retrograde flow of treadmilling actin filaments. Therefore, shootin1b KO reduces the impedance on the flow, resulting in an increase in the flow velocity (magenta arrow). This in turn reduces F-actin assembly translocation velocity (green arrow). Actin polymerization rate is not affected by shootin1b KO (Fig. 1*H*).

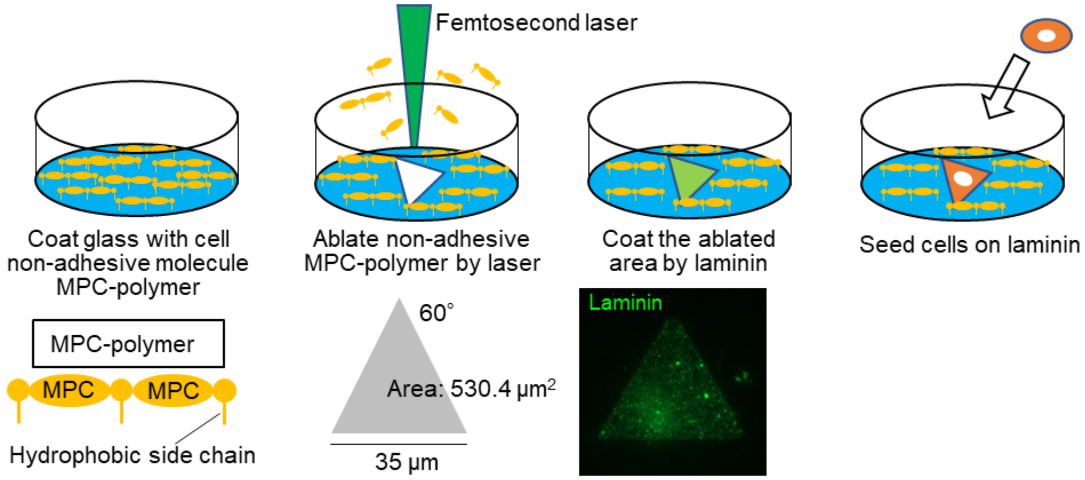


**Fig. S3.** Preparation of a laminin-coated adhesive island. A triangular pattern of adhesive laminin substrate with the edge length 35 μm and the area 530.4 μm$^2$ was created. The cell-adhesive patterns generated were confirmed by coating with green fluorescent-HiLyte 488-Laminin (for details see Materials and Methods).

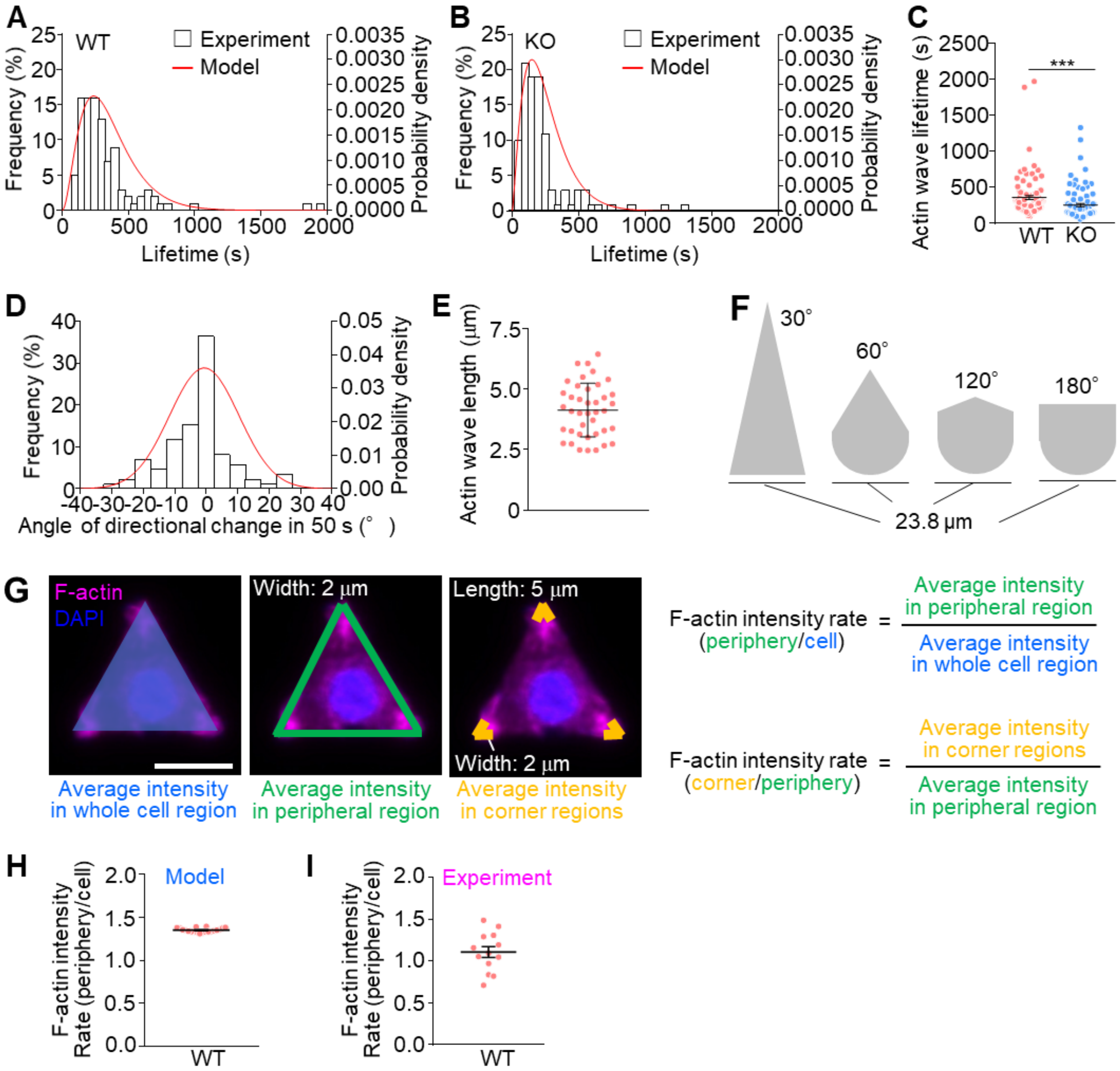


**Fig. S4.** (*A* and *B*) Distribution of the lifetime of filopodium-type SpTAs in WT (*A*) and shootin1b KO#1 U251 cells (*B*). White bars indicate the experimental data, while red curves show gamma distributions with order $\boldsymbol{\alpha}$ = 3.0 and $\boldsymbol{\beta}$ = 119 (A; WT), and with $\boldsymbol{\alpha}$ = 2.4 and $\boldsymbol{\beta}$ = 106 (*B*; shootin1b KO#1). The parameters were chosen to give the best fit to the experimental data by maximum likelihood estimation. (*C*) Lifetime of filopodium-type SpTAs in WT and shootin1b KO#1 U251 cells (WT, KO#1, N = 9 cells, n = 100 SpTAs). Data represent means ± SEM; ***p < 0.01. Statistical analysis was performed using the two-tailed Mann–Whitney *U*-test. (*D*) Distribution of the degree of the change in travelling direction of filopodium-type SpTAs in 50 sec. White bars indicate the experimental data, while red curves show Gaussian distributions with order mean; $\boldsymbol{\mu}$ = -0.61 and SD; $\boldsymbol{\sigma}$ = 11. The parameters were chosen to give the best fit to the experimental data by maximum likelihood estimation (N = 8 cells, n = 86 SpTAs). (*E*) Length of filopodium-type SpTAs (N = 4 cells, n = 40 SpTAs). (*F*) The pattern of adhesive laminin substrate with different corner angles, 30º, 60º, 120º or 180º, and the same width 23.8 µm and the same area 530.4 µm$^2$ used in the experiments in Fig. 3*G*. (*G*) The accumulation of actin filaments at the cell periphery was determined by dividing the average actin intensity in the peripheral region (green region; 2 µm width) by the average actin intensity in the whole cell region (blue region). Similarly, the accumulation at corners was determined by dividing the average of actin intensity in the corner regions (yellow regions; 2 µm width, 5 µm length) by the average actin intensity in the peripheral region (green region). Scale bar, 20 µm. (*H* and *I*) The accumulation rates of actin filaments at cell periphery in model data (H, n = 20) and U251 cells (I, N = 3 experiments, n = 13 cells). Data represent means ± SEM.

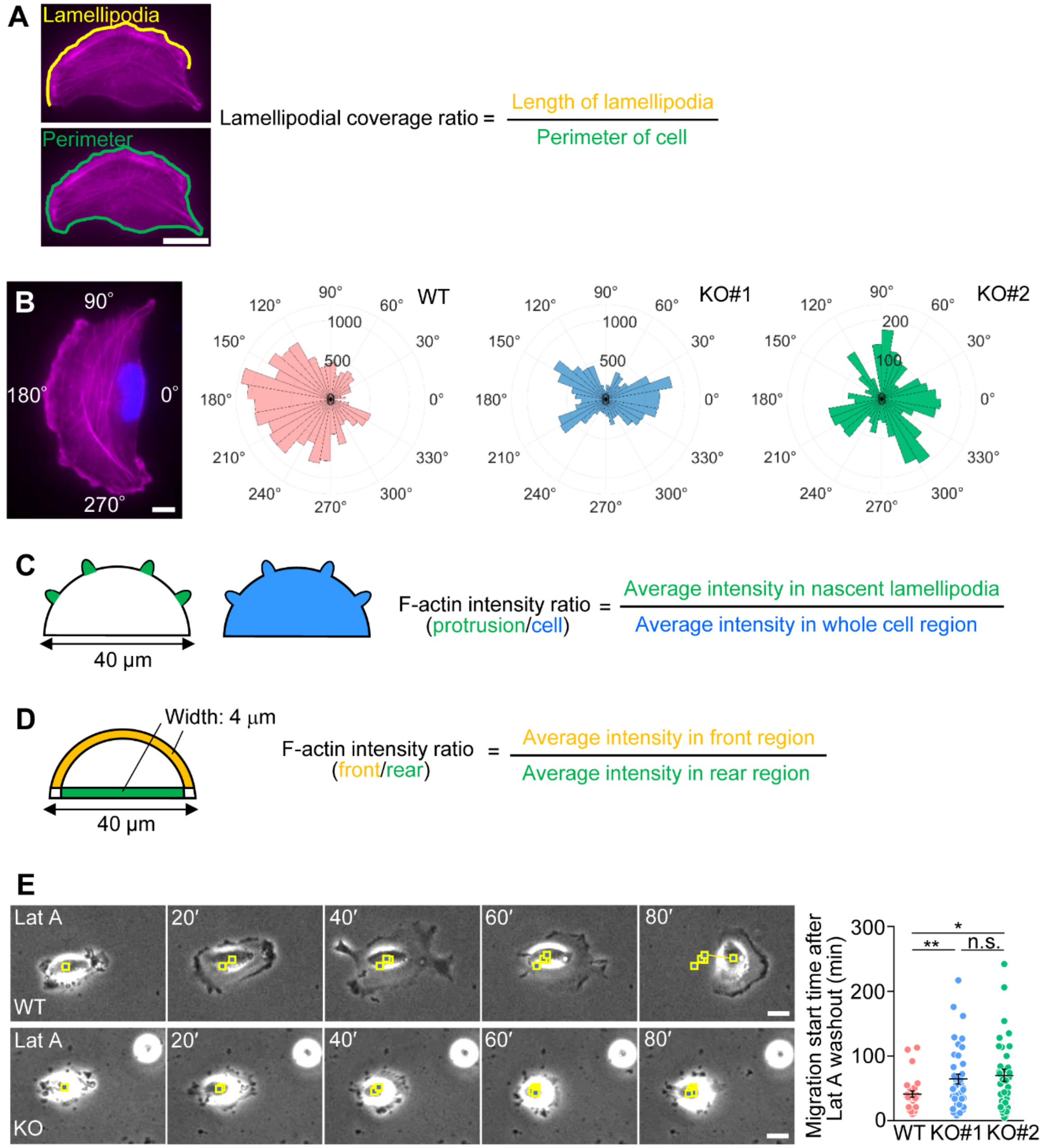


**Fig. S5.** (*A*) The lamellipodial coverage rate was determined by dividing the total length of lamellipodia (yellow line) by the perimeter of the cell (green line). Scale bar, 20 µm. (*B*) A WT U251 cell stained with Alexa Fluor 594 conjugated phalloidin and DAPI (left) and histogram of lamellipodia localization at cell periphery (right) of WT, shootin1b KO#1 and KO#2 U251 cells (WT, KO#1, KO#2, N = 3 experiments, n = 35 cells). Scale bar, 5 µm. Data represent means ± SEM; ***$p < 0.01$; *$p < 0.05$; ns, not significant. (*C*) The accumulation of actin filaments at the nascent lamellipodia was determined by dividing the average fluorescence intensity of actin filaments in protrusive regions (green regions) by the average intensity of actin filaments in the whole cell region (blue region). (*D*) Accumulation of actin filaments at the integrated lamellipodium was determined by dividing the average intensity of actin filaments at the convex leading edge (yellow region; 4 µm-width) by the average intensity of actin filaments in the rear region (green region; 4 μm-width). (*E*) Time-lapse phase-contrast images of WT and shootin1b KO#1 U251 cells taken after the treatment with 100 nM Lat A and at the indicated times after the Lat A washout. Yellow boxes trace cell movement. The right graph indicates the time required to start cell migration after the Lat A washout (WT, N = 3 experiments, n = 28 cells; KO#1, N = 3 experiments, n = 40 cells; KO#2, N = 3 experiments, n = 35 cells). Scale bar, 20 µm. Data represent means ± SEM; $^{**}p < 0.02$; $^{*}p < 0.05$; ns, not significant. Statistical analyses were performed using the two-tailed Mann–Whitney *U*-test. See also Fig. S4.

**Movie S1. Related to Figure 1**

First part: U251 cells expressing EGFP-LifeAct observed by TIRF microscopy. Left: filopodium-type and lamellipodium-type F-actin assemblies emerged widely in the cytoplasm, moved in random directions, and disappeared (see Fig. 1*A*). Right: filopodium-type F-actin assemblies changed the direction of their translocation randomly (yellow arrowheads); some emerged and separated (yellow and cyan arrowheads) from lamellipodium-type assemblies (arrow) or branched off (cyan arrowheads) from filopodium-type assemblies (see Fig. 1*B*). Some lamellipodium-type F-actin assemblies expanded from the filopodium-type assemblies (arrow, see Fig. S1). Time interval: 10 sec. Scale bar: 20 µm (left), 10 µm (right).

Second part: F-actin assemblies moving on the ventral plasma membrane of a COS7 cell expressing LifeAct-mCherry observed by TIRF microscopy (see Fig. S1*D*). Time interval: 60 sec. Scale bar: 20 µm.

**Movie S2. Related to Figure 1**

First part: an F-actin assembly translocating through the cytoplasm without interacting with the plasma membrane (see Fig. 1*D*). A U251 cell expressing LifeAct-mCherry was observed by 3D imaging with confocal deconvolution microscopy. Time interval: 13 sec. Z-stack interval: 0.2 µm. Scale bar: 5 µm.

Second part: a U251 cell expressing LifeAct-mCherry observed by TIRF microscopy and EGFP-LifeAct observed by epifluorescence microscopy (see Fig. S1*E* and *F*). F-actin assemblies observed by epifluorescence microscopy and TIRF microscopy are indicated by green and magenta arrowheads, respectively. Time interval: 60 sec.

**Movie S3. Related to Figure 1**

Fluorescent speckle dynamics of HaloTag-actin in an F-actin assembly; actin filaments were also monitored by EGFP-LifeAct. They were observed by TIRF microscopy. The F-actin assembly underwent treadmilling and translocated in the direction of polymerization (see Fig. 1*E*). A Time interval: 10 sec. Scale bar: 1 µm.

**Movie S4. Related to Figure 2**

Filopodia formation by filopodium-type F-actin assemblies (arrowheads) (see Fig. 2*A*). A U251 cell expressing EGFP-LifeAct was observed by epifluorescence microscopy. Time interval: 10 sec. Scale bar: 5 µm.

**Movie S5. Related to Figure 2**

Lamellipodium formation by a lamellipodium-type F-actin assembly (yellow arrow) (see Fig. 2*B*). A U251 cell expressing LifeAct-mCherry was observed by TIRF microscopy. The lamellipodium-type F-actin assembly merged with the pre-existing lamellipodia (cyan arrow) through lateral movement. Time interval: 60 sec. Scale bar: 5 µm.

**Movie S6. Related to Figure 2**

Filopodium-type F-actin assemblies entering into pre-existing lamellipodium (left) and filopodium (asterisk, right). The F-actin assembly entry resulted in local actin filament accumulation and lamellipodial expansion (arrow) (See Fig. 2*C*). U251 cells expressing EGFP-LifeAct were observed by TIRF microscopy. Time interval: 10 sec. Scale bars: 1 µm.

**Movie S7. Related to Figure 2**

Filopodium-type F-actin assemblies travelling along the lateral membranes (see Fig. 2*E*). U251 cells expressing LifeAct-mCherry were observed by epifluorescence microscopy. Time interval: 10 sec. Scale bars: 5 µm.

**Movie S8. Related to Figure 3**

Filopodium-type SpTAs (arrowheads) and a lamellipodium-type SpTA (arrow) accumulating at protrusive regions (asterisks) (see Fig. 3*A* and *B*). U251 cells expressing EGFP-LifeAct were observed by epifluorescence microscopy. Time interval: 10 sec. Scale bars: 5 µm (left), 20 µm (right).

**Movie S9. Related to Figure 3**

SpTA accumulation at the corners of a triangular U251 cell (see Fig. 3D). A U251 cell expressing LifeAct-mCherry was cultured on a triangular adhesive island and observed by epifluorescence microscopy. SpTAs travelled along the lateral edge, resulting in local accumulation of actin filaments at the corners upon arrival (arrowheads). Time interval: 300 sec. Scale bar: 10 μm.

**Movie S10. Related to Figure 3**

Mathematical model data showing SpTA accumulation in WT and shootin1b KO triangular cells (see Fig. 3*E* and *H*). Time interval: 10 sec. Scale bars: 5 μm.

**Movie S11. Related to Figure 4**

Fluorescence time-lapse Videos of WT and shootin1b KO#1 U251 cells expressing LifeAct-mCherry and observed by epifluorescence microscopy (see Fig. 4*C* and *F*). The images were taken before and after the treatment with 100 nM latrunculin A, and after the Lat A washout. Time interval: 60 sec. Scale bars: 20 μm.
